# Immunofocusing and enhancing autologous Tier-2 HIV-1 neutralization by displaying Env trimers on two-component protein nanoparticles

**DOI:** 10.1101/2020.11.30.403543

**Authors:** Philip J. M. Brouwer, Aleksandar Antanasijevic, Marlon de Gast, Joel D. Allen, Tom P. L. Bijl, Anila Yasmeen, Rashmi Ravichandran, Judith A. Burger, Gabriel Ozorowski, Jonathan L. Torres, Celia LaBranche, David C. Montefiori, Rajesh P. Ringe, Marit J. van Gils, John P. Moore, Per Johan Klasse, Max Crispin, Neil P. King, Andrew B. Ward, Rogier W. Sanders

## Abstract

The HIV-1 envelope glycoprotein trimer is poorly immunogenic because it is covered by a dense glycan shield. As a result, recombinant Env glycoproteins generally elicit inadequate antibody levels that neutralize clinically-relevant, neutralization-resistant (Tier-2) HIV-1 strains. Multivalent antigen presentation on nanoparticles is an established strategy to increase vaccine-driven immune responses. However, due to nanoparticle instability in vivo, the display of non-native Env structures, and the inaccessibility of many neutralizing antibody (NAb) epitopes, the effects of nanoparticle display have been modest for Env trimers. Here, we generated two-component self-assembling protein nanoparticles presenting twenty SOSIP trimers of the clade C Tier-2 genotype 16055. An immunization study in rabbits demonstrated that these nanoparticles induced 60-fold higher autologous Tier-2 NAb titers than the corresponding SOSIP trimers. Epitope mapping revealed that nanoparticle presentation focused antibody responses to an immunodominant apical epitope. Thus, these nanoparticles are a promising platform to improve the immunogenicity of Env trimers with apex-proximate NAb epitopes.

## Introduction

The immune evasion mechanisms and profound sequence diversity of human immunodeficiency virus-1 (HIV-1) make the development of a vaccine that can induce persistent and broad protection one of the greatest challenges in vaccinology today. To cope with the extensive diversity of globally circulating strains it is widely agreed that an effective vaccine will need to induce broadly neutralizing antibodies (bNAbs)^1–3^. Generally, bNAbs emerge after multiple years of virus-antibody coevolution in infected individuals; they typically recognize conserved epitopes on the densely glycosylated envelope glycoprotein (Env), the sole target of NAbs^4,5^. The development of recombinant immunogens that closely resemble the native trimeric conformation of Env, such as those based on the SOSIP design, have accelerated HIV-1 vaccine development by enabling the elicitation of autologous NAb responses in various animal models^2,6^. Further structure-guided engineering efforts have created trimers with improved stability, antigenicity and binding to germline B cells from various bNAb lineages^7–10^. Although consistent induction of bNAbs has so far not been achieved, several immunization regimens have been able to elicit antibodies with an encouraging although still inadequate degree of neutralization breadth^11,12^.

With most of their peptidic surface covered by glycans, native-like Env trimers are poorly immunogenic antigens^13^. Hence, several techniques have been developed in recent years to improve their ability to induce NAbs^14,15^. One strategy that has garnered increasing interest is multivalent antigen display. Presenting immunogens in a repetitive array on submicron particles has been shown to enhance critical immunological processes such as lymph node trafficking and B cell activation, leading to significantly increased neutralizing responses^16–18^. However, compared to the successes of particulate antigen display for RSV, HPV, and influenza^19–21^, the improvements to HIV-1 Env immunogenicity have been relatively modest^22^. Depending on the nanoparticle design, this may have been caused by factors such as nanoparticle instability in vivo, such as the uncoupling of trimer-liposome conjugations in vivo, or the heterogeneity of the displayed Env trimers, exemplified by the display of both non-native and native(-like) trimers on ferritin and virus-like particles^23–25^. In addition, the inaccessibility of multiple NAb epitopes on Env nanoparticles may further contribute to the modest effects that multivalent display confers to this particularly densely glycosylated glycoprotein^26,27^.

We recently described the design and characterization of two-component self-assembling protein nanoparticles presenting various SOSIP trimers^27–30^. These systems permit the purification of the trimer-bearing component prior to particle assembly in vitro, ensuring that exclusively native-like trimers are presented. Genetic fusion of SOSIP trimers to the trimeric nanoparticle subunit, I53-50A, enables the expression of SOSIP-I53-50A fusion proteins which, after mixing with a pentameric nanoparticle subunit, I53-50B.4PT1, assemble in vitro into monodisperse nanoparticles presenting twenty native-like SOSIP trimers (SOSIP-I53-50NPs). I53-50NPs presenting SOSIP trimers based on a consensus sequence of group M isolates (ConM SOSIP) were significantly better than the corresponding soluble trimers at inducing NAbs in immunized rabbits, but no benefit was seen when the I53-50NPs instead displayed SOSIP trimers of the BG505 genotype^27^. The inconsistent outcome was suggested to be attributable to differences in epitope accessibility on the two SOSIP trimers in the context of the nanoparticle surface. Whereas, ConM SOSIP trimers display an immunodominant NAb epitope near the trimer apex, which is highly accessible on the nanoparticles, the dominant NAb epitopes on BG505 SOSIP trimers are located much closer to the trimer base and are far less available. Thus, our findings suggested that SOSIP-I53-50NPs may be particularly suitable for SOSIP trimers that present an immunogenic neutralizing epitope near the apex^27^. A caveat is that ConM is a neutralization sensitive (Tier-1A) virus, so it remained uncertain whether the same scenario would apply to SOSIP trimers based on Tier-2 isolates.

Martinez-Murillo *et al.* described liposomes presenting native flexibly linked (NFL) Env trimers based on 16055; a clade C viral isolate with a Tier-2 phenotype^11,31^. These immunogens induced autologous NAb responses in rhesus macaques that were mapped to the apex-proximate V1/V2 region. Thus, unlike BG505, 16055 may be an appropriate Env genotype to investigate if SOSIP-I53-50NPs can improve the immunogenicity of SOSIP trimers based on a Tier-2 virus Env. Here, we describe the design, assembly, in vitro characterization and immunogenicity of 16055 SOSIP-I53-50NPs, and show that this nanoparticle platform can be beneficial to the immunogenicity of native-like Env trimers that display an apex-proximate NAb epitope.

## Results

### Improved trimerization, thermostability and antigenicity of 16055 SOSIP by introduction of a combination of stabilizing mutations

Our initial efforts to design a 16055 SOSIP trimer focused on the previously described SOSIP.v5.2 framework in which the A73C and A561C mutations introduce an extra disulfide bond between gp120 and gp41 (Fig. 1a and Supplementary Table 1)^9^. The resulting 16055 SOSIP.v5.2 construct was expressed in 293F cells and purified using PGT145 affinity chromatography. While this protein was fully cleaved, negative-stain electron microscopy (nsEM) showed a relatively high percentage of monomers and dimers, which is suggestive of instability (72% native-like trimers, 28% monomers/dimers; Fig. 1b and Supplementary Fig. 1). In an attempt to improve trimizeration and thermostability, and minimize the exposure of non-NAb epitopes, we combined several sets of mutations that have been shown to improve native-like trimers. These included the TD8 set of mutations^32^, the MD39 set^8^, and the T569G change^7^ (Fig. 1a and Supplementary Table 1). The resulting construct, from here on referred to as 16055 SOSIP.v8.3, was cleaved efficiently and visual inspection by nsEM revealed that >95% of the sample were now native-like trimers (Fig. 1b and Supplementary Fig. 1). Furthermore, nanoscale differential scanning fluorimetry (NanoDSF) demonstrated that these mutations collectively increased the midpoint of thermal denaturation (*T*m) by 13°C, from 66°C for 16055 SOSIP.v5.2 to 79°C for 16055 SOSIP.v8.3 (Fig. 1c).

**Fig. 1.**
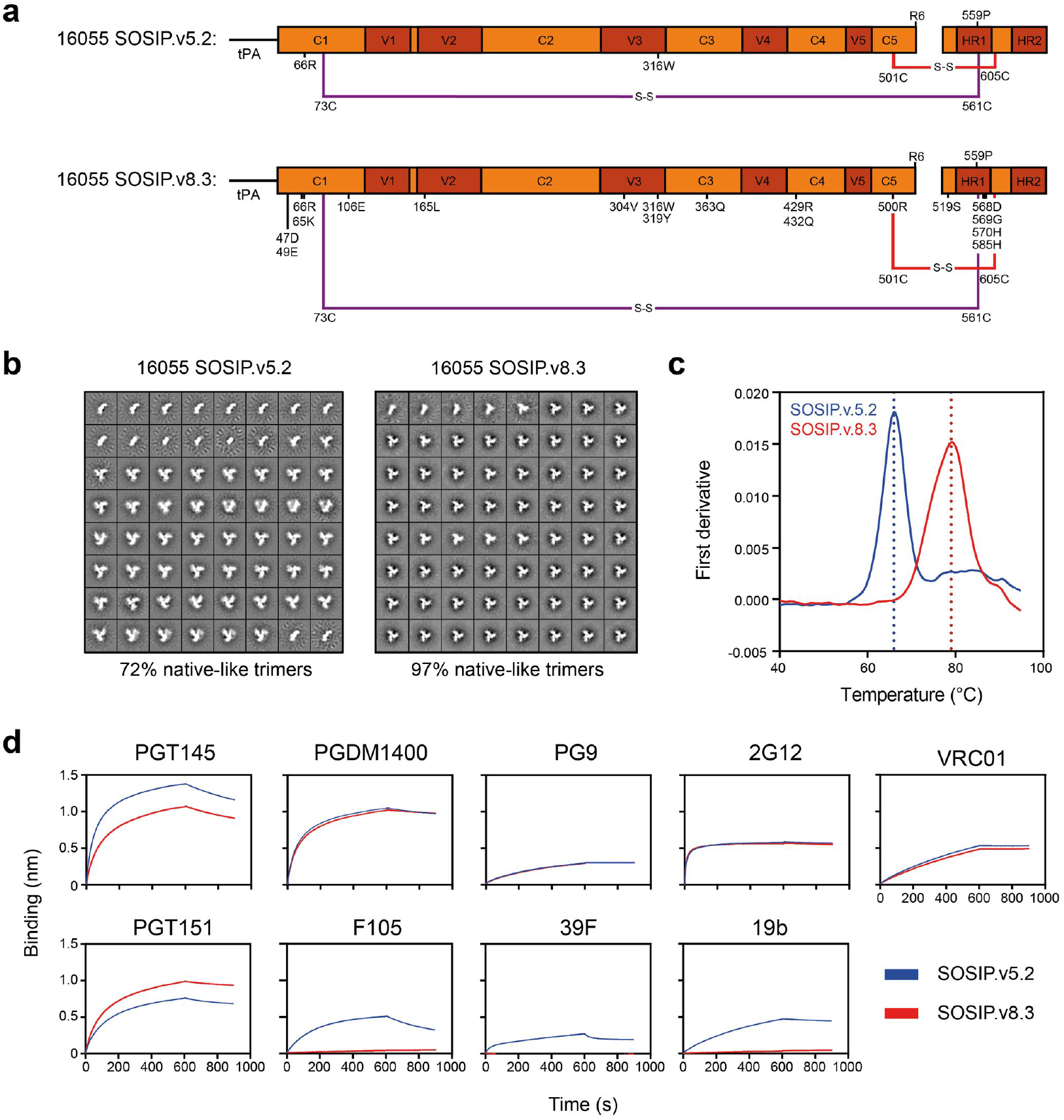
Biophysical and antigenic characterization of 16055 SOSIP.v5.2 and 16055 SOSIP.v8.3. **a** Linear schematic of the 16055 SOSIP.v5.2 and 16055 SOSIP.v8.3 constructs with mutations annotated. The two disulfide bonds that link gp120 and gp41 are shown in red or purple. tPA = tissue plasminogen activator signal peptide. **b** 2D-class averages from nsEM from 16055 SOSIP.v5.2 (left) and 16055 SOSIP.v8.3 (right) with the percentages of native-like trimers shown below. **c** Thermostability of 16055 SOSIP.v5.2 and 16055 SOSIP.v8.3 as determined by NanoDSF. The dotted lines indicate the corresponding *T*m. Shown are representative melting curves from three technical replicates. **d** Sensorgrams from BLI experiments showing binding of 16055 SOSIP.v5.2 and 16055 SOSIP.v8.3 to immobilized bNAbs (PGT145, PGDM1400, PG9, 2G12, VRC01, and PGT151) and non-NAbs (F105, 39F, and 19b).

To assess whether 16055 SOSIP.v8.3 had an improved antigenic profile, we performed bio-layer interferometry (BLI) experiments to analyze the binding of both trimer variants to various immobilized bNAbs and non-NAbs. The bNAbs PG9, PGDM1400, 2G12, and VRC01 bound to both trimers with indistinguishable kinetics (Fig. 1d). The quaternary-dependent bNAb PGT151 bound slightly better to SOSIP.v8.3, while the opposite was seen with PGT145. Furthermore, whereas 16055 SOSIP.v5.2 bound detectably to the CD4bs-directed non-NAb F105 and V3-directed non-NAbs 19b, and 39F, such binding was completely abrogated for 16055 SOSIP.v8.3 (Fig. 1d). Thus, the introduction of SOSIP.v8.3 mutations markedly improved the trimer’s antigenic profile by greatly reducing the exposure of non-NAb epitopes. The 16055 SOSIP.v8.3 construct, hereafter referred to as 16055 SOSIP for convenience, was used in all subsequent experiments.

### Efficient assembly of I53-50NPs presenting 16055 SOSIP trimers

To produce SOSIP-I53-50NPs presenting 16055 SOSIP trimers, we first genetically fused 16055 SOSIP to the trimeric I53-50A component as described previously^27^. The resulting construct could be produced to reasonable yields (0.7 mg/L) after expression in 293F cells and PGT145-purification and was fully cleaved as demonstrated by reducing SDS-PAGE (Supplementary Fig. 1). To form SOSIP-I53-50NPs, PGT145-purified 16055 SOSIP-I53-50A trimers were further purified by size exclusion chromatography (SEC) and mixed with I53-50B.4PT1 in an equimolar ratio, followed by an overnight incubation at 4°C and a second SEC step to remove any unassembled components (Fig. 2a). SDS-PAGE and nsEM analyses of the pooled fractions demonstrated that the SEC-purified 16055 SOSIP-I53-50NPs consisted of well-ordered icosahedral nanoparticles that displayed fully cleaved SOSIP trimers (Fig. 2b and Supplementary Fig. 1). A very small population of aggregated nanoparticles was observed by nsEM. This was confirmed by dynamic light scattering experiments that showed, in addition to a monodisperse population of particles with the expected hydrodynamic radius, a small percentage of larger particles (Supplementary Fig. 2.). Consistent with previous data on other trimer genotypes, the assembly of 16055 SOSIP-I53-50NPs was highly efficient, as only trace amounts of unassembled components were visible in the SEC fractions. Thermal melting experiments by NanoDSF showed that 16055 SOSIP-I53-50NPs displayed a typical two-phase melting profile with an unfolding peak at 76°C for the SOSIP trimers and a second peak at 87°C for the I53-50NP core (Supplementary Fig. 2). To further gauge the stability of 16055 SOSIP-I53-50NPs we incubated them at 37°C in phosphate-buffered saline (PBS) for up to 48 hours, and performed a native PAGE assay. Particles remained intact over the duration of this experiment (Supplementary Fig. 2).

**Fig. 2.**
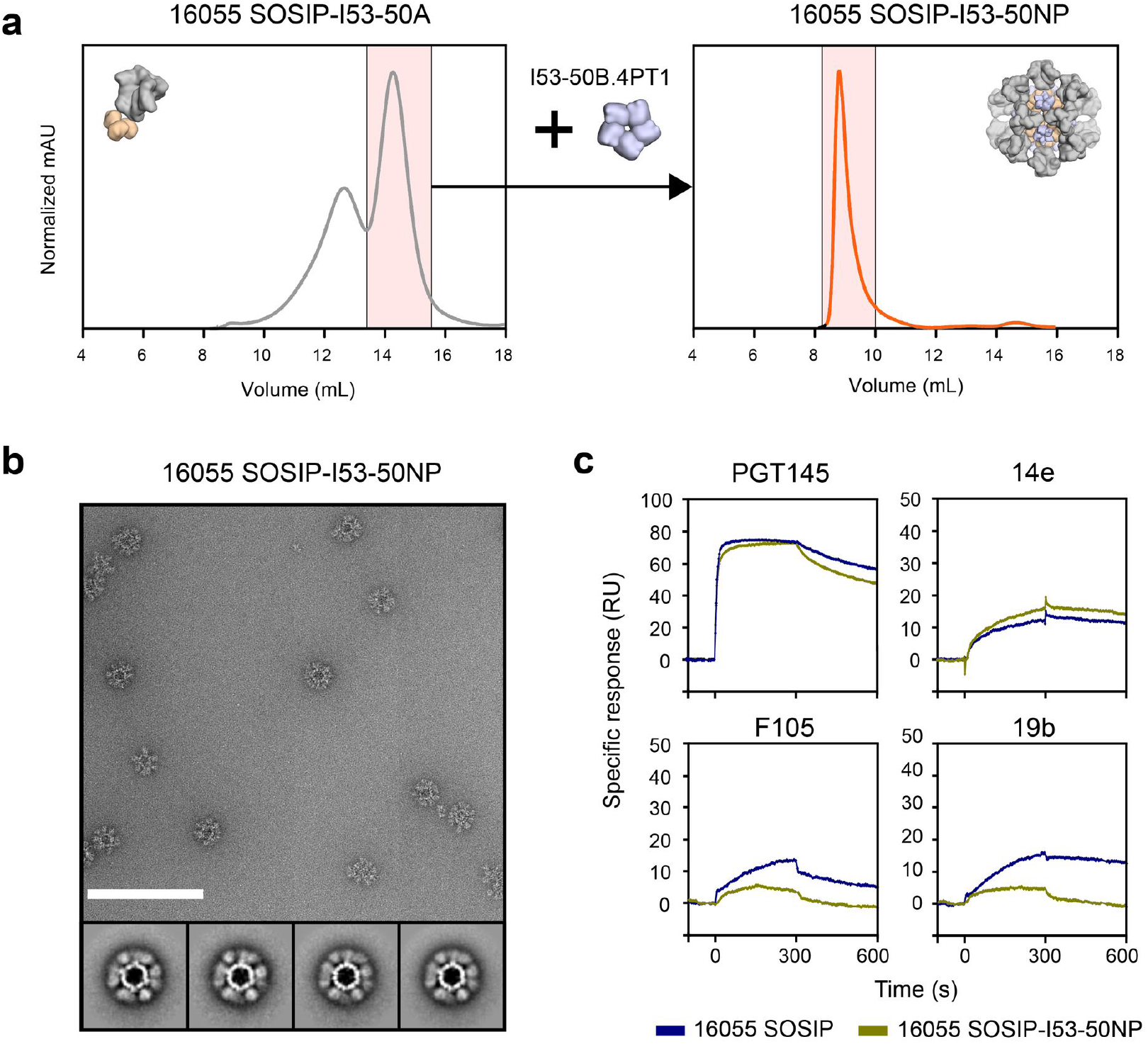
Biophysical and antigenic characterization of 16055 SOSIP-I53-50NPs. **a** Representative size exclusion chromatograph of 16055 SOSIP-I53-50A (left panel) and the assembled 16055 SOSIP-I53-50NP (right panel). The fractions of 16055 SOSIP-I53-50A that were collected and mixed with I53-50B.4PT1 (left panel), and the fractions that were collected to purify the assembled 16055 SOSIP-I53-50NPs (right panel), are shown in pink shading. **b** Raw nsEM image of the SEC-purified 16055 SOSIP-I53-50NPs with 2D-class averages shown below. White scale bar corresponds to 200 nm. **c** Sensorgrams from SPR experiments showing the binding of bNAb PGT145 and non-NAbs 14e, F105 and 19b to either 16055 SOSIP or 16055 SOSIP-I53-50NPs, as color coded in the legend below.

Next, we performed surface plasmon resonance (SPR) experiments to assess whether 16055 SOSIP trimers maintained their structural integrity on the nanoparticles. 16055 SOSIP trimers and corresponding nanoparticles were immobilized onto chips by their His-tags (for SOSIP trimers on the C terminus, for SOSIP-I53-50NPs on the exterior-facing C terminus of the I53-50B.4PT1 pentamer) and the binding of a panel of monoclonal antibodies (mAbs) was monitored (for more details see the Methods section). Nanoparticles showed similar binding of the trimer-dependent bNAb PGT145, although with marginally faster dissociation from the trimers displayed on I53-50NPs. The non-NAb 14e bound only slightly better to nanoparticles than the corresponding trimers, while binding by 19b and F105 was strongly reduced for nanoparticles. The latter suggests that linkage of 16055 SOSIP to I53-50A or its assembly into nanoparticles slightly improves its antigenicity. Overall, we concluded that presentation of 16055 SOSIP trimers on I53-50NPs had no adverse impact on trimer antigenicity (Fig. 2c).

### Comparative glycan analysis of 16055 SOSIP and 16055 SOSIP-I53-50A

To determine the site-specific compositions and occupancy of glycans on the 16055 SOSIP and the corresponding nanoparticle, we performed nanoliquid chromatography-electrospray ionization-mass spectrometry (nanoLC-ESI MS) (Fig. 3 and Supplementary Table 2). Glycopeptides of sufficient quality could be obtained for all potential N-linked glycosylation sites (PNGS) except N392 and N399 for 16055 SOSIP and N230, N392, and N399 for 16055 SOSIP-I53-50A. First, we assessed whether particular PNGS were occupied by a glycan and to what extent. We observed several underoccupied sites on the gp120 subunit of 16055 SOSIP. Specifically, N156 and N160 were occupied by a glycan on only 64% and 67% of the trimers, respectively, while N136, N197, and N230 were also underoccupied (i.e. <90% occupancy). All other PNGS on gp120 were fully occupied (>90% occupancy). On the gp41 subunit, N611 and N616 were underoccupied, with an occupancy of 71% and 40%, respectively. This is consistent with previous observations with other SOSIP trimers, and possibly a consequence of their close proximity to each other and/or the presence of serines instead of threonines at the +2 position^33^. N625 and N637 were fully occupied on 16055 SOSIP. Overall, PNGS were occupied similarly on 16055 SOSIP-I53-50A compared to SOSIP, with some exceptions: N145, N156, N301, and N409 were less occupied on SOSIP-I53-50A, while N136, N160, N197, N611, and N616 were occupied more often.

**Fig. 3.**
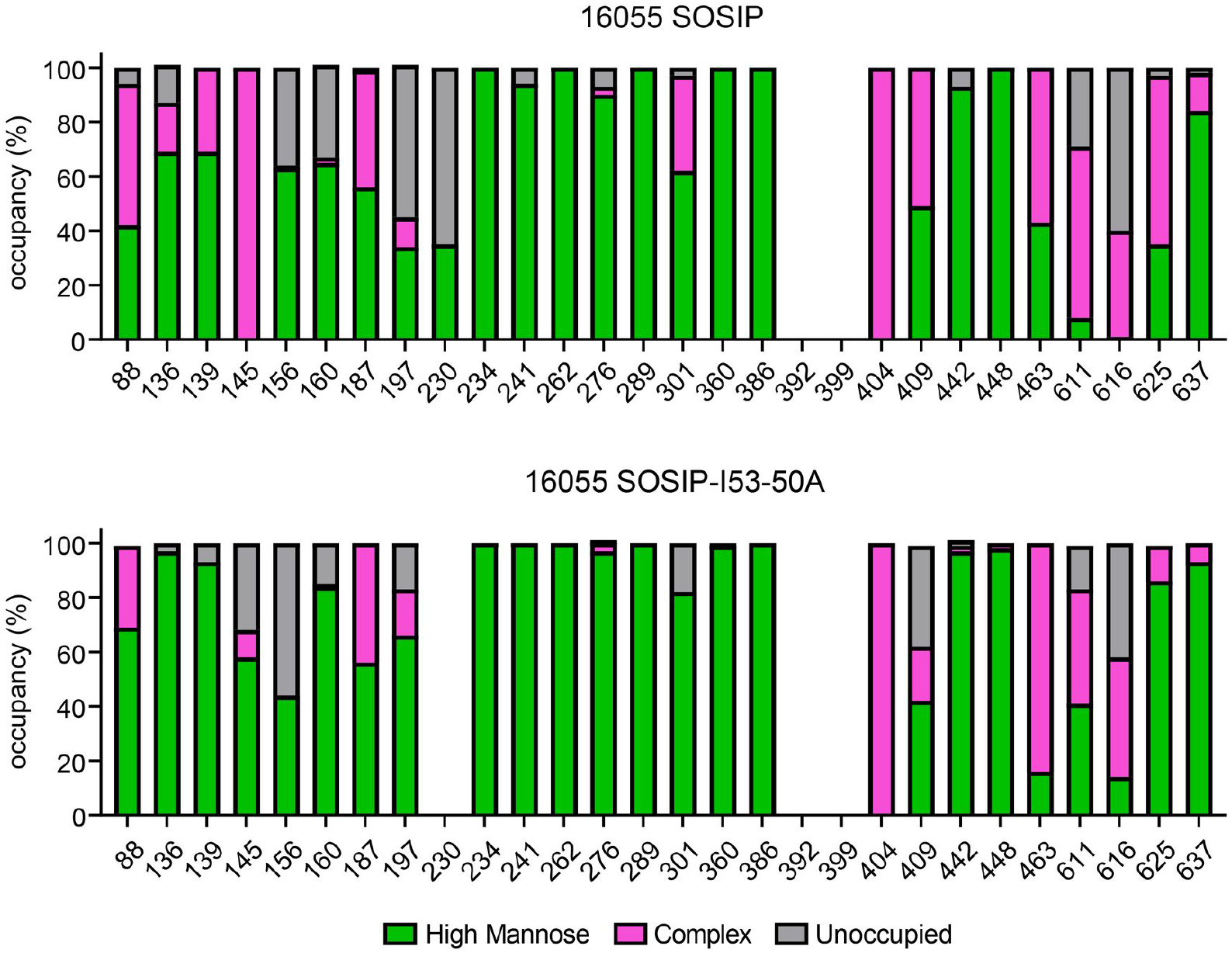
Site specific glycosylation of 16055 SOSIP and 16055 SOSIP-I53-50A. Quantification of site specific occupancy and composition for all 28 glycan sites on 16055 SOSIP trimers (top) and 16055 SOSIP-I53-50A trimers (bottom) derived from nanoLC-ESI MS.

As with SOSIP trimers of other genotypes, the glycans on gp120 of 16055 SOSIP were mostly of the oligomannose type, while complex glycans predominated on gp41 (Fig. 3 and Supplementary Table 2)^34,35^. The glycan composition of the previously described 16055 NFL trimer was very similar to that of 16055 SOSIP, with differences limited to N301 (oligomannose glycans on 16055 NFL), and N136, N139, N187, N409, and N463 (complex glycans on 16055 NFL)^11^. The addition of I53-50A component at the C-terminus of 16055 SOSIP, required for nanoparticle formation, appears to have markedly restricted glycan processing at sites N88, N136, N139, N145, N301, N611, and N625 (Fig. 3 and Supplementary Table 2). At other sites the glycan compositions were very similar on the standard and I53-50A versions of the 16055 SOSIP trimer. We thus conclude that 16055 SOSIP-I53-50A had a glycan shield that was not dramatically different from that of 16055 SOSIP.

### Epitope-dependent enhancement of SOSIP-specific B cell activation in vitro

The high antigen density on nanoparticles can enhance the activation of cognate B cells by allowing high avidity interactions with B cell receptors (BCRs), but steric constraints can also reduce epitope accessibility. To study this key topic, we generated B cells expressing BCRs for various bNAbs and measured the ability of 16055 SOSIP-I53-50NPs to trigger Ca^2+^ influx in vitro. The B cells expressing the apex-targeting bNAbs PGDM1400 and VRC26 were efficiently activated by 16055 SOSIP-I53-50NPs, but the corresponding 16055 SOSIP trimers were ineffective (Fig. 4). A similar outcome was seen with B cells bearing BCRs for the VRC01 CD4-binding site bNAb. Soluble trimers did activate B cells expressing the V3-glycan bNAb PGT121, but SOSIP-I53-50NPs did so markedly more efficiently.

**Fig. 4.**
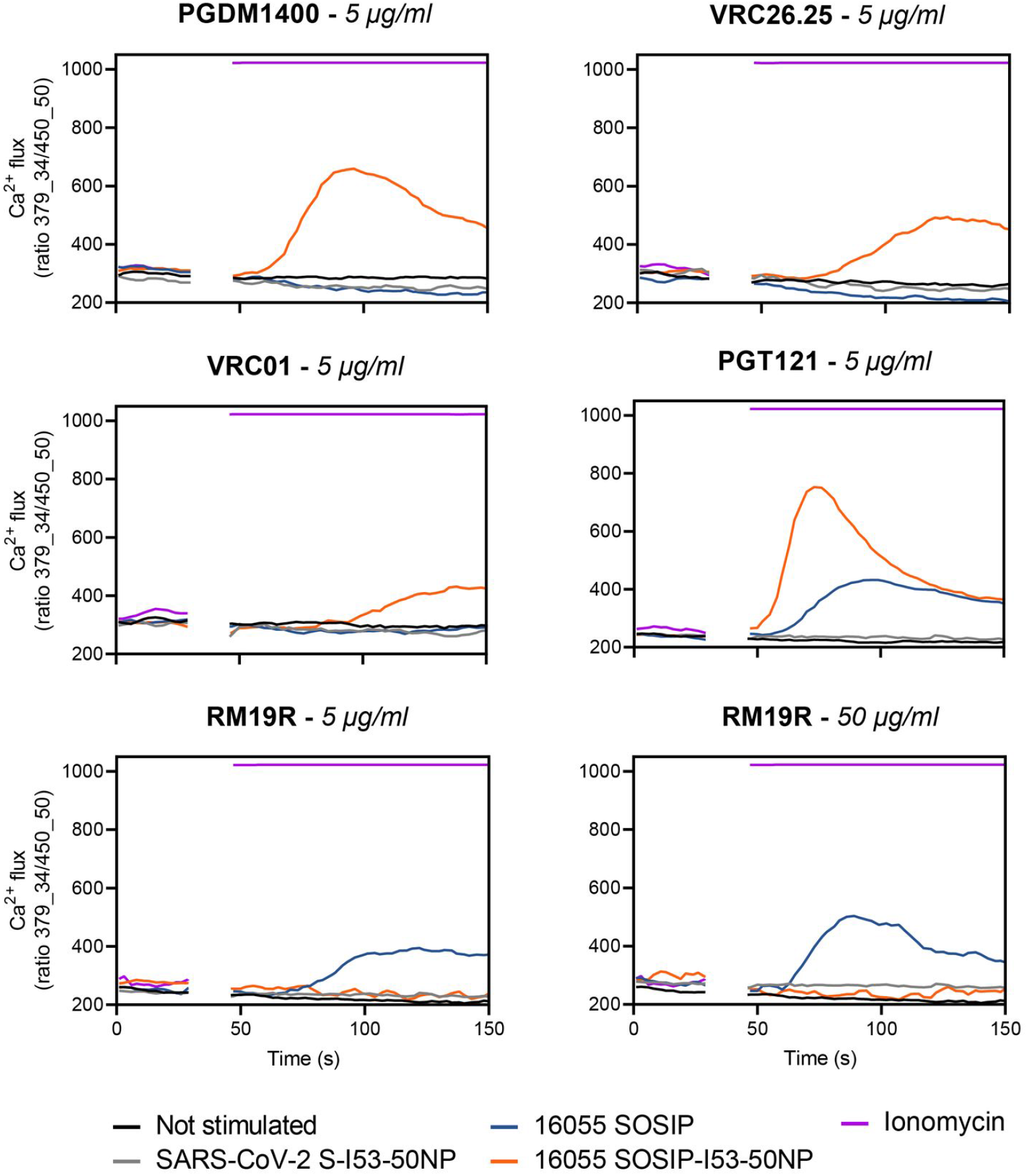
B cell activation analysis of 16055 SOSIP and 16055 SOSIP-I53-50NPs. Activation of B cells expressing PGDM1400, VRC26.25, PGT121, VRC01 or RM19R was assessed by measuring Ca^2+^ influx into the cells. After 30 seconds of baseline measurement, an equimolar amount of 16055 SOSIP either as a soluble antigen or presented on I53-50NPs was added to the B cells and Ca^2+^ flux was measured over ~100 seconds. The concentration of soluble 16055 SOSIP that was used in each experiment is indicated in italics in each graph title. Ionomycin and SARS-CoV-2 S-I53-50NP served as positive and negative controls, respectively.

We also generated B cells expressing the base-specific non-NAb RM19R that was isolated from a BG505 SOSIP trimer-immunized macaque^36^. The exposed base of SOSIP trimers contains immunodominant non-NAb epitopes that may distract the immune response away from sites more favored to NAb induction^37^. While soluble 16055 SOSIP trimers strongly activated B cells bearing RM19R BCRs, this was not seen when 16055 SOSIP-I53-50NPs were tested, even at 10-fold higher concentration (Fig. 4). In control experiments, I53-50NPs expressing SARS-CoV-2 Spike proteins did not activate any of the B cell lines. We conclude that presenting 16055 SOSIP trimers on I53-50NPs increases the efficiency of B cell activation in an epitope-dependent manner. Apex-proximate bNAb epitopes benefit from the higher antigen density present on the NPs, but in contrast a non-NAb epitope is now inaccessible to its BCR.

### Increased autologous neutralization by presentation of 16055 SOSIP on I53-50NPs

To assess whether presentation on I53-50NPs increased the immunogenicity of 16055 SOSIP trimers, New Zealand White rabbits (n=5 per group) were immunized at weeks 0, 4 and 20 with 30 μg of SOSIP trimer or the equimolar amount presented on nanoparticles, formulated in the adjuvant Adjuplex. Rabbits were bled at various time points and immunogenicity was assessed using both anti-trimer ELISAs and neutralization assays that included autologous and heterologous pseudoviruses (Fig. 5a).

**Fig. 5.**
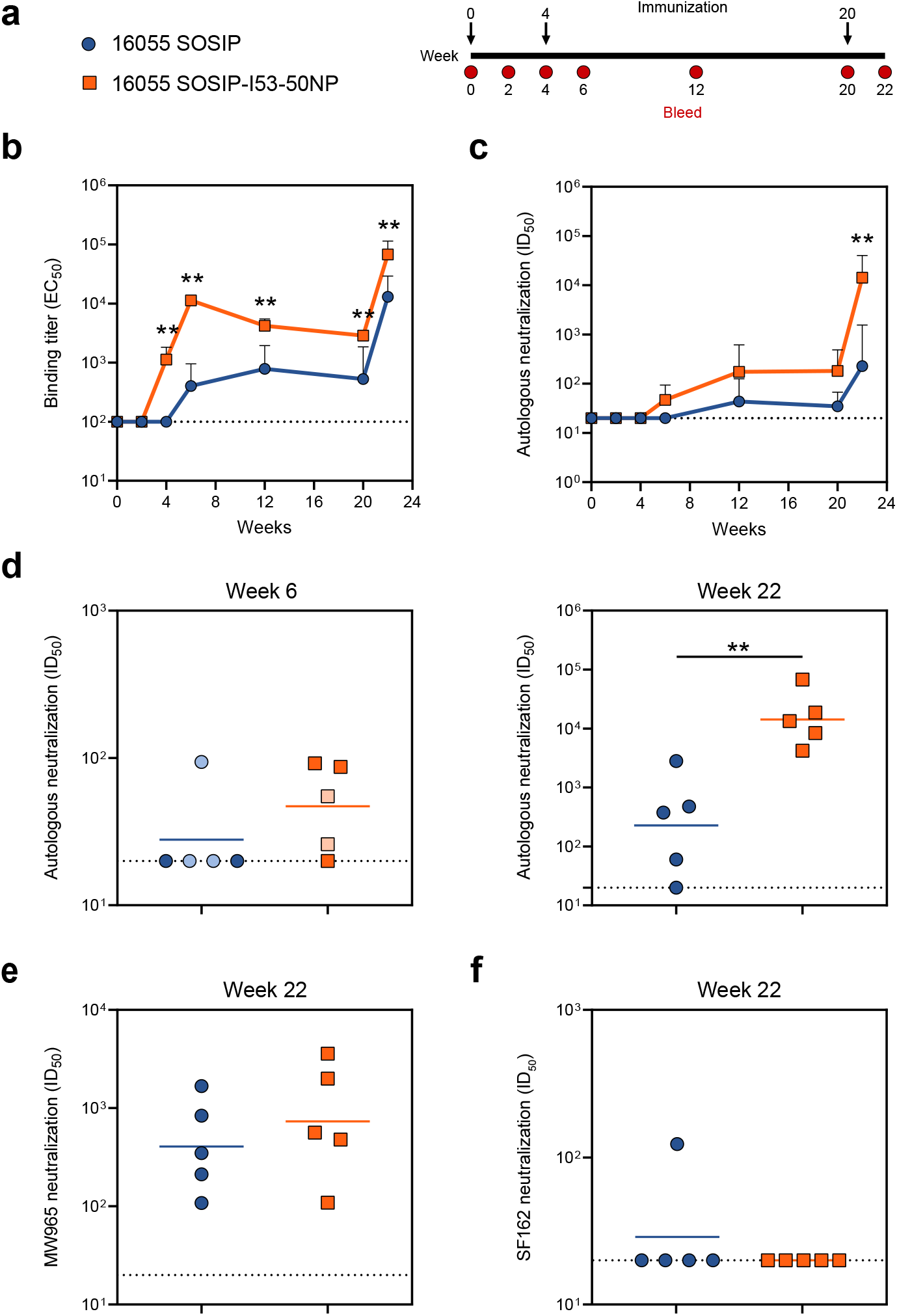
Immunogenicity of 16055 SOSIP trimers and 16055-I53-50NPs in rabbits. **a** Schematic representation of the immunization schedule with color coding for each immunogen. **b-f** Statistical differences between two groups (*n* = 5 rabbits) were determined using two-tailed Mann–Whitney *U*-tests (\**p* < 0.05; \*\**p* < 0.01). Dotted lines indicate the lowest serum dilution. **b** Midpoint binding titers against 16055 SOSIP over time. Shown are geometric means with standard deviations. **c**Midpoint 16055 pseudovirus neutralization titers over time. Shown are geometric means with standard deviations. **d** Midpoint 16055 pseudovirus neutralization titers at week 6 (left panel), and week 22 (right panel). Symbols with lighter tints represent rabbit sera that had a high MLV background (ID50,16055 is less than 3x ID50,MLV). Horizontal bars indicate geometric mean values. See also Supplementary Table 3 for replicate data of week 22, performed at Duke University Medical Center. **e** Midpoint MW965 pseudovirus neutralization titers at week 22. Horizontal bars indicate the geometric mean. **f** SF162 midpoint neutralization titers at week 22. Horizontal bars indicate geometric mean values.

16055 SOSIP induced binding antibody titers that were detectable after the second immunization at week 6 and were further boosted by the third immunization. From week 4 onwards, i.e. after only one immunization, 16055 SOSIP-I53-50NP consistently induced significantly higher binding titers against 16055 SOSIP than the corresponding trimer (two-tailed Mann-Whitney *U*-test; week 4, week 6, week 12, week 20, and week 22: *p*=0.0079) (Fig. 5b). As expected, 16055 SOSIP-I53-50NPs also induced a detectable antibody response against the I53-50NP cage (Supplementary Fig. 2). Whereas 16055 SOSIP trimers did not induce any 16055 pseudovirus neutralization titers above cut-off two weeks after the prime and first boost (except for one rabbit that had a high non-specific NAb titer against MLV), 2/5 rabbits that received 16055 SOSIP-I53-50NP developed modest NAb responses following the second immunization (Fig. 5c and d). The largest difference between the two groups was observed at week 22, after the third immunization, with nanoparticles inducing a 60-fold higher NAb titer than corresponding trimers (geometric mean NAb titer of 14272 versus 227, respectively; two-tailed Mann-Whitney *U*-test; *p*=0.0079) (Fig. 5d and Supplementary Table 3). NAb titers were also higher for the 16055 SOSIP-I53-50NP recipients, at weeks 12 and 20, although this was not statistically significant.

In addition to autologous neutralization, we analyzed the sera’s ability to neutralize heterologous Tier-1 and Tier-2 pseudoviruses. Neutralization of the former is often indicative of the elicitation of non-NAbs that target the V3-loop; an epitope which is accessible on non-native Env forms and perhaps when native-like trimers become destabilized (in vivo)^38,39^. Sera from trimer and nanoparticle recipients neutralized the Tier-1 pseudovirus MW965 with similar potencies (Fig. 5e and Supplementary Table 3). The SF162 Tier-1 pseudovirus was, however, at most minimally neutralized, possibly related to structural differences in the V3 loop of 16055 and SF162 (Fig. 5f). To assess if 16055 SOSIP-I53-50NPs were also able to broaden the NAb response, neutralization of a panel of 9 Tier-2 pseudoviruses representing the global HIV-1 diversity was tested. However, sera from both groups of rabbits neutralized these heterologous pseudoviruses only sporadically and to very low levels (Supplementary Table 4).

### Immunofocussing on an apex-proximate NAb epitope by presentation of 16055 SOSIP on I53-50NPs

Our previous work has suggested that presentation on I53-50NPs is particularly beneficial for the immunogenicity of SOSIP trimers with an apex-proximate immunodominant NAb epitope. To visualize the specificities of the polyclonal antibody response after immunization with either 16055 SOSIP trimers or 16055 SOSIP-I53-50NPs, we performed electron microscopy-based polyclonal epitope mapping (EMPEM). Fabs from week 22 serum were complexed with 16055 SOSIP trimers and imaged using nsEM (Fig. 6a and Supplementary Fig. 3).

**Fig. 6.**
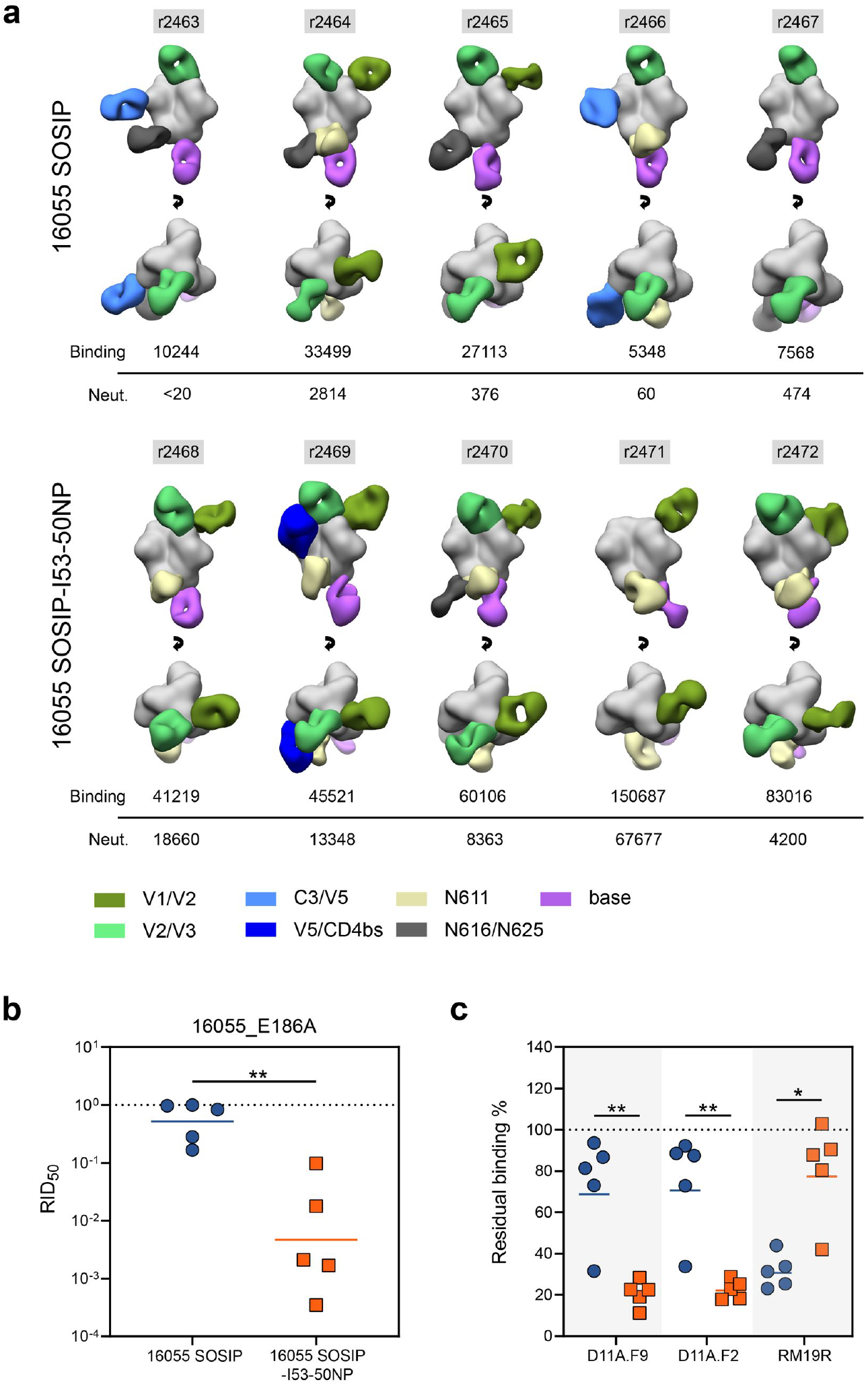
Epitope mapping of 16055 SOSIP and 16055 SOSIP-I53-50NP-induced antibody responses. **a** Composite figures showing binding of polyclonal Fabs from week 22 sera to 16055 SOSIP trimers as determined by EMPEM analysis. The rabbit ID’s are indicated above each figure, and autologous midpoint binding and neutralization titers are shown below. The Fabs are color coded based on their apparent epitopes as indicated in the legend at the bottom. Only a single Fab is depicted for each epitope cluster for simplicity. **b** RID50 of the 16055_E186A pseudovirus mutant relative to the parental 16055 pseudovirus for 16055 SOSIP and 16055 SOSIP-I53-50NP. The dotted line indicates a RID50 of 1, i.e. the E186A substitution does not affect neutralization. **c** Residual binding of D11A.F9, D11A.F2 and RM19R after incubation of sera with 16055 SOSIP in ELISA. The dotted line indicates 100% residual binding, i.e. the serum does not compete with binding of the mAb. **b, c** Symbols represent individual rabbit sera as shown in figure 4. Horizontal bars indicate the geometric mean values. Statistical differences were determined using two-tailed Mann–Whitney *U*-tests (\**p*<0.05; \*\**p* < 0.01).

The EMPEM analyses revealed seven distinct regions that were targeted by antibodies: two sites overlapping with variable loops on gp120 (V1/V2 and V2/V3), two other sites on gp120 (C3/V5 and V5/CD4bs), two sites on gp41 centered on PNGSs (N611 and N616/N625), and finally a site involving the trimer base. Rabbits immunized with 16055 SOSIP-I53-50NPs showed a consistent V1/V2-directed response (5/5 rabbits) in contrast to the trimer recipients (2/5 rabbits). These antibodies showed a very similar angle of approach and epitope location as the 16055-specific neutralizing mAbs which were isolated from macaques immunized with 16055 NFL trimers^31^. The V1/V2 Ab specificities observed by EMPEM may, therefore, be neutralizing. Indeed, with the exception of rabbit 2467, the presence of these responses tracked with neutralization (Fig. 6a).

To confirm that these V1/V2-targeting antibodies correlated with neutralization, we performed a neutralization assay with a 16055 pseudovirus containing an E186A substitution in V1. This substitution was previously shown to abrogate neutralization by macaque-derived 16055-specific NAbs^31^. Introduction of an alanine at position 186 drastically decreased neutralization of the 16055 pseudovirus by 6/7 of the rabbit sera that showed V1/V2-responses by EMPEM (Fig. 6b). With a geometric mean relative NAb titer (RID50, the ratio of NAb titers of the 16055 E186A pseudovirus mutant relative to the parental 16055 pseudovirus) of ~0.005 (i.e. a ~200-fold decrease in NAb titers by introduction of E186A) the effect of the substitution was particularly marked for the nanoparticle group. Furthermore, we observed a significant correlation between the RID50, which is a proxy for V1/V2-directed NAb responses, and 16055 neutralization titers (Spearman-rank correlation; *p*=<0.0001, *r*=−0.976) (Supplementary Fig. 4). This provided further evidence that the V1/V2 apex is the immunodominant neutralizing epitope on 16055 SOSIP trimers.

To provide a more quantitative analysis of the V1/V2-directed antibody response we performed a competition ELISA with two 16055-specific NAbs that target the V1/V2 apex: D11A.F2 and D11A.F9^31^. Sera from the trimer group competed significantly less efficiently with these two mAbs than those from the nanoparticle group (two-tailed Mann-Whitney *U*-test; D11A.F2 and D11.F9: *p=*0.0079) (Fig. 6c). In addition, competition of D11A.F2 and D11A.F9 correlated significantly with autologous neutralization, which is additional evidence that neutralization is predominantly a result of antibodies targeting the V1/V2 epitope (Supplementary Fig. 4). We also performed a competition ELISA with the base-specific and non-NAb RM19R. While EMPEM showed base-specific responses in all the 16055 SOSIP-I53-50NP recipients (Fig. 6a), sera from 16055 SOSIP-I53-50NP rabbits competed significantly less efficiently with RM19R than the trimer recipients (Fig. 6c and Supplementary Fig. 4). Thus, although 16055 SOSIP-I53-50NPs induce base-specific antibody responses, they do so significantly less efficiently compared to 16055 SOSIP trimers.

In addition to base-specific responses, we observed several other antibody responses by EMPEM which showed little correlation with neutralization as they were present in both neutralizing and non-neutralizing serum samples. All but one rabbit (r2471) developed antibody responses that targeted an epitope that overlapped the V2/V3 loop around glycans at position 156 and 160 (Fig. 6a). Considering the large glycan under-occupancy at these sites and the fact that breaches in the glycan shield generally create immunogenic epitopes^34,40^, it is conceivable that these antibodies were elicited as a result of partial glycosylation (Fig. 3). Similarly, glycan under-occupancy may have resulted in the elicitation of antibodies targeting the base-proximate N616/N625 epitope as well as the N611 epitope. The higher frequency of N616/N625 responses in sera from the trimer recipients may be explained by the lower occupancy at the N616 PNGS on 16055 SOSIP compared to 16055 SOSIP-I53-50A (Fig. 3). Antibody responses targeting the C3/V5-loops (r2463 and r2466) and V5/CD4bs (r2469) were unique to the trimer and nanoparticle groups, respectively.

## Discussion

The development of two-component protein nanoparticles that self-assemble in vitro has allowed efficient, scalable and controlled production of nanoparticle immunogens for several class I viral fusion proteins^19,27,30,41^. While presenting RSV-F or influenza HA on these nanoparticles increased vaccine-elicited NAb responses by at least 10-fold, the effects of nanoparticle display for HIV-1 Env have not yielded similarly positive results^19,41^. In the context of the trimer’s dense glycan shield, the effect of multivalent display for HIV-1 Env is influenced by the location and accessibility of its NAb epitope(s)^26,27^. We previously showed that presenting a SOSIP trimer on I53-50 nanoparticles can increase the magnitude of the NAb response when the immunodominant NAb epitope is located near the apex of the trimer^27^. To confirm and extend these previous results using a SOSIP trimer based on a neutralization-resistant Tier-2 pseudovirus, we generated SOSIP-I53-50NPs based on the 16055 genotype. When the re-engineered and stabilized 16055 SOSIP trimers were presented on I53-50NPs, there was a significant increase in NAb responses, compared to soluble trimers, that were focused on a previously described V1/V2-apex epitope^31^.

After three immunizations 16055 SOSIP-I53-50NPs induced 60-fold higher NAb titers than the corresponding trimers. For comparison, after three immunizations, liposomes presenting noncovalently linked NFL 16055 Env trimers induced a 3-fold higher geometric mean NAb titer over its soluble trimeric counterpart^31^. While there are differences in the adjuvant, the immunization schedule and the animal model between that study and ours, the extent of the difference is still notable. One reason for the apparently greater benefit of the SOSIP-I53-50NP design may be superior stability under in vivo conditions, which is reportedly problematic for liposomes with noncovalently attached Env trimers^24^. The more uniform size and/or the well-ordered and potentially more optimal geometry of the SOSIP-I53-50NP's may also be relevant.

Even though it is hard to pinpoint exactly which immunological processes underlie the marked improvements in NAb titers conferred by I53-50NP display, our findings suggest an important role for immunofocussing. This is supported by the observed higher frequency of V1/V2-specific NAbs in the EMPEM analysis and the competition ELISA which showed that presentation of 16055 SOSIP on I53-50NPs shifts the antibody response from base- to apex- dominated specificities. Together, these results are in line with our findings that nanoparticle display particularly enhances the activation of B cells that are specific for apex-proximate epitopes. Although we did not specifically address this, it is conceivable that the higher NAb titers in the nanoparticle group are a result of improved affinity maturation of the antibody response as multiple studies have shown that multivalent antigens enhance processes critical for affinity maturation such as follicular dendritic cell deposition^42–45^.

While our findings imply that the increased NAb titers by 16055 SOSIP-I53-50NPs are a direct effect of multivalent presentation, we cannot rule out that subtle differences in glycosylation between the two immunogens have also contributed to the altered immunogenicity. Site-specific glycan analysis revealed several glycan sites that were partially occupied on 16055 SOSIP. The responses targeting the area overlapping the V2/V3 loop, the N611 PNGS, and the N616/N625 PNGS might have developed as a consequence of underoccupancy at positions N160/N156, N611, and N616/N625, respectively. Restoring glycan occupancy at these sites, such as by altering PNG sites from NxS to NxT^33^, may focus responses away from these non-NAb epitopes toward NAb-relevant regions such as the V1/V2-loop or the base-proximate bNAb epitope near glycan N88^11^. Furthermore, considering the role of glycans N160 and N156 in PGT145 binding, restoring occupancy of these sites may also improve the 16055 SOSIP yields when PGT145 is used as the purification method^46^.

The competition ELISA with RM19R showed that the non-neutralizing base epitope was significantly less immunogenic when 16055 SOSIP was presented on I53-50NPs. However, base-specific antibody responses were still present in 16055 SOSIP-I53-50NP recipients, as shown by EMPEM analysis and the low levels of serum antibody competition with RM19R. This result was unexpected considering that the accessibility of these epitopes is greatly reduced on the assembled nanoparticle^27^, and we showed here that 16055 SOSIP-I53-50NPs cannot activate in vitro B cells bearing BCRs for base epitope non-NAbs (Fig. 4). Thus, it is therefore likely that the anti-base antibodies arose from nanoparticle disassembly in vivo. Additional efforts to better understand and overcome any stability limitations to SOSIP-I53-50NPs are justified and in progress.

In contrast to ConM SOSIP-I53-50NPs, where the improvement in NAb titers waned during subsequent boosts^27^, 16055 SOSIP-I53-50NPs demonstrated the greatest improvement over soluble trimer after the final immunization. The varying outcomes may reflect differences in the intrinsic immunogenicity of the two SOSIP trimer genotypes. For example, the dominant (Tier-1) NAb epitope on soluble ConM SOSIP trimers may be sufficiently immunogenic that there is eventually no benefit to nanoparticle presentation once the responses maximize over time. For the 16055 SOSIP trimer, with a less immunogenic (Tier-2) NAb epitope, nanoparticle display may still be advantageous even after several immunizations. Recent relevant findings imply that multivalent presentation is particularly useful for immunogens that have a low affinity for cognate B cells^47^. More immunization studies with SOSIP-I53-50NPs of different genotypes could allow us to more fully understand the dynamics of how NAb responses to different epitopes are generated by immunogens of varying valencies.

Although autologous NAb titers were improved the 16055 SOSIP-I53-50NPs did not induce broader responses. This is not surprising in light of how the HIV-1 vaccine field has come to understand the requirements for inducing true bNAb-like antibodies. Nonetheless, I53-50NPs remain a promising nanoparticle platform as they significantly increase the immunogenicity of dominant and apex-proximate NAb epitopes. Using SOSIP-I53-50NPs to multivalently present germline targeting trimers or implementing them in sequential immunogen regimens^10,11,48^ is an obvious next step to optimize vaccination strategies that aim to induce HIV-1 bNAbs.

## Materials and Methods

For further information and material requests please contact Rogier Sanders

### Construct design

The untagged 16055 SOSIP.v5.2 construct was generated by digesting a pPPI4 plasmid with PstI and NotI and replacing the previously described BG505 SOSIP.v5.2 with an ordered 16055 SOSIP sequence (Integrated DNA Technologies) that had the furin cleavage site (REKR) substituted for RRRRRR and the following mutations introduced: H66R, A73C, T316W, A501C, I559P, A561C, and T605C. To create the 16055 SOSIP.v8.3-I53-50A construct, a modified 16055 SOSIP.v5.2 sequence was ordered (Integrated DNA Technologies) and cloned by Gibson assembly into the previously described I53-50A plasmid which was digested using PstI and BamHI^27^. Modification consisted of the following previously described mutations: E47D, K49E, V65K, I165L, E429R, R432Q, A500R (The TD8 mutations^32^), R304V, A319Y, S363Q, I519S, L568D, V570H, R585H (the MD39 mutations^8^), and T569G^7^. To create the untagged 16055 SOSIP.v8.3 construct, a stop-codon was introduced directly downstream of the C-terminal 664 residue. For Ni-NTA ELISAs and SPR experiments, His-tagged versions of 16055 SOSIP.v8.3 were generated by addition of a GSGSGGSGHHHHHHHH amino acid sequence immediately after the C-terminal residue 664 of the trimer.

### Env protein expression and purification

All 16055 constructs were expressed in transiently transfected HEK293F cells (Invitrogen, cat no. R79009) maintained in Freestyle medium (Life Technologies) using previously described methods^38,49^. Briefly, a 3:1 mixture of PEImax (1 μg/μL; 937.5 μg/L cells) and expression plasmids (312.5 μg/L cells) were added to 0.8-1.2 million cells/mL. To ensure optimal furin-mediated cleavage of the Env component, the cells were transfected with (1) a 4:1 ratio of Env plasmid and furin for SOSIP constructs or (2) a 3:1 ratio of Env plasmid and furin for SOSIP-I53-50A constructs. Proteins were purified from vacuum-filtered (0.22 μm filters) transfection supernatants by PGT145 bNAb-affinity chromatography. PGT145-coupled sepharose beads were mixed with the filtered supernatant and after an overnight incubation on a roller, subjected to washing and elution as described earlier^49^. Protein eluates were concentrated and buffer exchanged to TN75 (75 mM NaCl, 20 mM Tris HCl pH 8.0) using Vivaspin filters with a 100 kDa molecular weight cutoff (GE Healthcare). Protein concentrations were determined using the Nanodrop method. The required proteins peptidic molecular weight and extinction coefficient were obtained by filling in the protein's amino acid sequence in the online Expasy software (ProtParam tool).

### I53-50B.4PT1 and I53-50A protein expression and purification

The I53-50A and I53-50B.4.PT1 proteins were expressed as described recently^45^. Briefly, Lemo21 cells(DE3) (NEB), which were grown in LB (10 g Tryptone, 5 g Yeast Extract, 10 g NaCl) in 2 L baffled shake flasks or a 10 L BioFlo 320 Fermenter (Eppendorf) were used to express the I53-50A or I53-50B.4PT1 proteins grown. Cells were induced with 1 mM IPTG and maintained shaken for ~16 h at 18°C. Microfluidization was used to harvest and lyse the cells, using a Microfluidics M110P machine at 18,000 psi in 50 mM Tris, 500 mM NaCl, 30 mM imidazole, 1 mM PMSF, 0.75% CHAPS. Proteins were purified by applying clarified lysates to a 2.6×10 cm Ni Sepharose 6 FF column (Cytiva) on an AKTA Avant150 FPLC system (Cytiva). A linear gradient of 30 mM to 500 mM imidazole in 50 mM Tris, pH 8, 500 mM NaCl, 0.75% CHAPS was used to elute both proteins. Next, the pooled fractions were subjected to size-exclusion chromatography on a Superdex 200 Increase 10/300, or HiLoad S200 pg GL SEC column (Cytiva) in 50 mM Tris pH 8, 500 mM NaCl, 0.75% CHAPS buffer. I53-50A elutes at ~0.6 column volume (CV) whereas I53-50B.4PT1 elutes at ~0.45 CV. Prior to nanoparticle assembly, protein preparations were tested to confirm low levels of endotoxin.

### I53-50NP assembly

To generate naked I53-50NP particles, a final concentration of 50 μM purified I53-50A and 50 μM I53-50B.4PT1 in 25 mM Tris pH 8, 500 mM NaCl, 0.75% CHAPS were incubated at RT for at least 1 hour while gently rocking. Next, the assembly mix was sterile filtered (0.22 μm) and applied to a Superose 6 increase 10/300 GL column (GE Healthcare) to remove unassembled components. Well-ordered nanoparticles eluted at a volume of 11.5 mL.

### SOSIP-I53-50NP assembly

16055 SOSIP-I53-50NPs were produced as described recently^27^. Briefly, PGT145-purified 16055 SOSIP-I53-50A was subjected to size-exclusion chromatography using a Superose 6 Increase (GE Healthcare) column in Assembly Buffer II (25 mM Tris, 500 mM NaCl, 5% glycerol, pH 8.2) to remove aggregated proteins. Column fractions that contained no aggregated trimers were then immediately pooled and mixed in an equimolar ratio with I53-50B.4PT1 followed by an overnight incubation step at 4°C. The assembly mix was concentrated at 350 × g using Vivaspin filters (10 kDa molecular weight cutoff; GE Healthcare) and subjected to another round of size-exclusion chromatography using the same column and buffer to remove unassembled components. Fractions corresponding to assembled nanoparticles (elution between 8.5-10.5 mL with a peak at 9 mL) were pooled and concentrated at 350 × g by spin filtration with Vivaspin filters (10 kDa molecular weight cutoff; GE Healthcare). 16055 SOSIP-I53-50NPs were then buffer exchanged into PBS supplemented with 250 mM sucrose using a Slide-A-Lyzer MINI dialysis device (20 kDa molecular weight cutoff; ThermoFisher Scientific). Nanoparticle concentrations were determined by the Nanodrop method using the particles peptidic molecular weight and extinction coefficient.

### SDS-PAGE and BN-PAGE analysis

SDS-PAGE and BN-PAGE were performed as described previously^38^. Briefly, for SDS-PAGE, 2 μg of SOSIP trimer or 3.2 μg of SOSIP-I53-50NP (which is equivalent to 2 μg of trimer) were loaded on a 4-12% Tris-Glycine gel or 6-18% Tris-Glycine gel (both from Invitrogen). For BN-PAGE, the same amounts of SOSIP trimer and SOSIP-I53-50NP were loaded on a 4-12% Bis-Tris NuPAGE gel or 3-12% Bis-Tris NuPAGE gel (both from Invitrogen), respectively.

### Bio-layer Interferometry

A concentration of 10 μg/mL of mAb in running buffer (PBS, 0.02% Tween20, 0.1% BSA) was loaded on ProtA biosensors using an Octet K2 (ForteBio). After dipping the chip in running buffer to remove excess mAb, the chip was dipped for 600 s in a well containing 100 μM of 16055 SOSIP.v5.2 or 16055 SOSIP.v8.3 in running buffer to measure association. Finally, the chip was dipped for 300 s in running buffer to measure dissociation.

### Differential Scanning fluorimetry

Prometheus NT.48 NanoDSF instrument (NanoTemper Technologies) was used for all the *T*m determination experiments, as described previously^27^. SOSIP trimer and SOSIP-I53-50NP samples (in triplicates) were diluted to 0.35 mg/mL in TBS and loaded into NanoDSF capillaries. The temperature was increased from 20 – 95°C at a rate of 1 °C/min. Instrument software was used to calculate the 1st derivative curve and the location of the maximum was taken as the *T*m value.

### Surface plasmon resonance

Surface Plasmon Resonance (SPR) was used for assessing the structural integrity of the 16055 SOSIP.v8.3 trimers on the I53-50NP surface, as described elsewhere^27^. All experiments were carried out on a BIAcore 3000 instrument (Cytiva, formerly GE Healthcare) at 25°C with HBS-EP (0.01 M HEPES, 0.15 M NaCl, 3 mM EDTA, 0.005% v/v Surfactant P20, pH 7.4) as running buffer.

C1 sensor chips were used for immobilizing His-tagged trimer and nanoparticles by anti-histidine antibody (Cytiva). Briefly, the C1 sensor surface was pre-washed and anti-histidine antibody (diluted to 50 μg/ml in Na-acetate pH 4.5) was amine-coupled in all flow cells, following the standard-amine coupling protocol, yielding densities of ~1500 RU. Trimers and nanoparticles were immobilized in parallel flow cells, at a mean density of 224 ± 0.862 RU (s.e.m.) and 316 ± 0.961 RU to give the same amounts of Env (i.e., to control for the extra non-Env mass in the nanoparticles, see also^27^). One flow cell served as a background control for non-specific binding. For binding analysis, 1 μM of each mAb was allowed to associate for 300 s and dissociate for 300 s, at a flow rate of 30 μl/min. At the end of each binding cycle, the sensor surface was regenerated using a single pulse of 10 mM Glycine (pH 2.0), at a flow rate of 75 μl/min. Each mAb was analyzed in duplicate cycles. The sensograms are presented after subtraction of non-specific binding in the reference flow cell (which was <1%).

### Site-specific glycan analysis using mass spectrometry

Env proteins were denatured for 1h in 50 mM Tris/HCl, pH 8.0 containing 6M of urea and 5 mM dithiothreitol (DTT). The Env proteins were reduced and alkylated by adding 20 mM iodacetamide (IAA) and incubated for 1h in the dark, followed by a 1h incubation with 20 mM DTT to eliminate residual IAA. The alkylated Env proteins were buffer-exchanged into 50 mM Tris/HCl, pH 8.0 using Vivaspin columns (3 kDa) and digested separately overnight using trypsin or chymotrypsin (Mass Spectrometry Grade, Promega) at a ratio of 1:30 (w/w). Peptides were dried and extracted using C18 Zip-tip (MerckMilipore). The peptides were dried again, re-suspended in 0.1% formic acid and analyzed by nanoliquid chromatography-electrospray ionization-mass spectrometry (LC-ESI MS) with an Easy-nLC 1200 (Thermo Fisher Scientific) system coupled to a Fusion mass spectrometer (Thermo Fisher Scientific) using higher energy collision-induced dissociation (HCD) fragmentation. Peptides were separated using an EasySpray PepMap RSLC C18 column (75 μm × 75 cm). The liquid chromatography conditions were as follows: 275-minute linear gradient consisting of 0-32% acetonitrile in 0.1% formic acid over 240 minutes followed by 35 minutes of 80% acetonitrile in 0.1% formic acid. The flow rate was set to 200 nL/min. The spray voltage was set to 2.7 kV and the temperature of the heated capillary was set to 40 °C. The ion transfer tube temperature was set to 275 °C. The scan range was 400-1600 m/z. The HCD collision energy was set to 50%, appropriate for fragmentation of glycopeptide ions. Precursor and fragment detection were performed using an Orbitrap at a resolution MS1= 100,000. MS2= 30,000. The AGC target for MS1=4e5 and MS2=5e4 and injection time: MS1=50ms MS2=54ms

Glycopeptide fragmentation data were extracted from the raw file using Byos (Version 3.9; Protein Metrics Inc.). The glycopeptide fragmentation data were evaluated manually for each glycopeptide; the peptide was scored as true-positive when the correct b and y fragment ions were observed along with oxonium ions corresponding to the glycan identified. The relative amounts of each glycan at each site as well as the unoccupied proportion were determined by comparing the extracted chromatographic areas for different glycotypes with an identical peptide sequence. The precursor mass tolerance was set at 4ppm for MS1 and 10ppm for fragments MS2. A 1% false discovery rate was applied. The relative amounts of each glycan at each site as well as the unoccupied proportion were determined by comparing the extracted ion chromatographic areas for different glycopeptides with an identical peptide sequence.

To obtain data for sites that frequently present low intensity glycopeptide the glycans present on the glycopeptides were homogenized to boost the intensity of these peptides. This analysis loses fine processing information but enables the ratio of oligomannose: complex: unoccupied to be determined. The remaining glycopeptides were first digested with Endo H (New England Biolabs) to deplete oligomannose- and hybrid-type glycans and leave a single GlcNAc residue at the corresponding site. The reaction mixture was then dried completely and resuspended in a mixture containing 50 mM ammonium bicarbonate and PNGase F (New England Biolabs) using only H2O^18^ (Sigma-Aldrich) throughout. This second reaction cleaves the remaining complex-type glycans but leaves the GlcNAc residues remaining after Endo H cleavage intact.

The use of H2O^18^ in this reaction enables complex glycan sites to be differentiated from unoccupied glycan sites as the hydrolysis of the glycosidic bond by PNGaseF leaves a heavy oxygen isotope on the resulting aspartic acid residue. The resultant peptides were purified as outlined above and subjected to reverse-phase nanoLC-MS. Instead of the extensive N-glycan library used above, two modifications were searched for: +203 Da corresponding to a single GlcNAc, a remnant of an oligomannose/hybrid glycan, and +3 Da corresponding to the O^18^ deamidation product of a complex glycan. A lower HCD energy of 27% was used as glycan fragmentation was not required. Data analysis was performed as above and the relative amounts of each glycoform determined, including unoccupied peptides. Data from the two sets of experiments was combined and glycans grouped according to the presence of high-mannose glycans, corresponding to hybrid- and oligomannose-type glycans, complex-type glycans and unoccupied PNGS.

### Negative Stain EM

Negative stain electron microscopy (nsEM) experiments were performed as previously described^27,28^. All samples (free trimers or assembled nanoparticles) were diluted to 20-50 μg/mL and loaded onto the carbon-coated Cu grids (400-mesh, glow-discharged at 15 mA for 30 s). The sample was blotted off with a filter paper and the grids were then negatively stained with 2 % (w/v) uranyl-formate for 60 s. Data was collected on a Tecnai F20 electron microscope, operating at 200 keV. Nominal magnification was 62,000 × with a resulting pixel size of 1.77 Å at the specimen plane. The defocus was −1.50 μm. Tietz 4k × 4k TemCam-F416 CMOS camera was used for imaging. Data collection was performed using Leginon automated imaging interface^50^. Initial data processing was performed using the Appion data processing suite^51^. For SOSIP-I53-50NP samples, approximately ~1,000 particles were manually picked from the micrographs and subjected to Iterative MSA/MRA algorithm for 2D classification. For free 16055 SOSIP trimers ~20,000 particles were auto-picked and 2D-classified.

### Generation of PGDM1400, VRC26.25, PGT121, VRC01 and RM19R-expressing B cells

To generate B cells stably expressing Env-specific BCRs, IgM-negative Ramos B cells^52^ were essentially subjected to lentiviral transduction. Lentiviruses were produced by co-transfecting a T25 flask of HEK 293T cells with the plasmids pMDL (1.5 μg), pVSV-g (0.83 μg) and pRSV-Rev (0.6 μg), and the BCR-encoding B-cell specific plasmid pRRL EuB29 gl2-1261 IgGTM.BCR.GFP.WPRE^53^ (2.4 μg) using lipofectamine 2000 (Invitrogen). Prior to lentiviral production, the gl2-1261 gene was exchanged by Gibson assembly with ordered heavy and light chain genes of either mature PGDM1400, VRC01, PGT121, VRC26.25 and RM19R (Integrated DNA Technologies). Two days post-transfection, supernatants containing lentiviral particles were filtered (0.45 μm) and concentrated to 200 μl using Vivaspin filters using Vivaspin filters with a 100 kDa molecular weight cutoff (GE Healthcare). Subsequently, IgM-negative Ramos B cells^52^ cultured in RPMI supplemented with 10% fetal calf serum, penicillin (100 U mL−1) and streptomycin (100 μg mL−1) were transduced with the concentrated viral supernatant. Seven days post-transduction, GFP and IgG double-positive cells (i.e. BCR-expressing cells) were sorted on a FACS ARIA-II SORP (BD Biosciences) and cultured.

### B cell activation experiments

B cell activation experiments of Ramos B cells stably expressing VRC01, VRC26.25, PGDM1400, PGT121 or RM19R BCRs were performed as previously described^27,49^. Briefly, BCR-expressing Ramos B cells were suspended at 4 million cells/mL in RPMI++ and labeled with 1.5 μM Indo-1 (Invitrogen) for 30 min at 37 °C. Cells were then washed with Hank’s Balance Salt Solution containing 2 mM CaCl2 and incubated for another 30 min at 37 °C. B cell Ca^2+^ influx was monitored by measuring the 379/450 nm emission ratio on a LSR Fortessa (BD Biosciences). Following 30 s of baseline measurement, aliquots of 1 million cells/mL were then stimulated for 100 s at RT, with 5 or 50 μg/mL of 16055 SOSIP or the equimolar amount presented on I53-50NPs. To determine the maximum Ca^2+^ influx, 1mg/mL ionomycin (Invitrogen) was used. Kinetics analyses were performed using FlowJo v10.7.

### Rabbit immunizations

Female and naive New Zealand White rabbits (2.5 – 3kg), were arbitrarily distributed among groups (5 rabbits per group) and were sourced and housed at Covance Research Products Inc. (Denver, PA, USA). Immunizations were performed under permits with approval number C0038-19. Rabbits recieved an intramuscular immunization in each quadricep at weeks 0, 4, and 20. Immunization mixtures consisted of 30 μg of 16055 SOSIP.v8.3 trimers or the equimolar amount presented on I53-50NPs (48 μg) formulated in Adjuplex adjuvant (Empirion). Calculations of the dose were based on the peptidic molecular weight of the proteins, which were obtained by filling in the protein's amino acid sequence in the online Expasy software (ProtParam tool). The rabbits were bled on the day of immunization and at weeks 2, 6, 12, and 22.

### IgG isolation and Fab purification

To purify polyclonal rabbit IgG, 1 mL of rabbit serum was incubated overnight with 250 μL of protein G agarose beads (ThermoScientific). The next day, IgGs were eluted using 0.1 M glycine, pH 2.5 after which the elution buffer was neutralized with 2M Tris, pH 8.0. Following concentration and buffer exchange to PBS with Vivaspin filters (10 kDa molecular weight cutoff; GE Healthcare), IgG concentrations were measured using the Nanodrop method. Next, to generate Fabs, purified IgG was incubated, shaking, with immobilized papain resin (ThermoScientific; 100 μL resin/mg of IgG) for 5 hrs at 37°C in phosphate buffer saline, 10 mM EDTA, 20 mM cysteine, pH 7.4. After 5 hrs the resin was removed by spin centrifugation using Spin-X^®^ centrifuge tube filters (Corning Inc.). The flow-through was then incubated with Protein A agarose resin (ThermoScientific; 200 μL resin/mg of Ig) for 2 hrs, shaking at RT to remove Fc and non-digested IgGs. After removing the resin by spin centrifugation using Spin-X^®^ centrifuge tube filters (Corning Inc.), the flow-through was buffer exchanged to PBS and concentrated using Vivaspin filters (10 kDa molecular weight cutoff; GE Healthcare).

### EMPEM analysis

Purified, polyclonal Fab samples were complexed with 16055 SOSIP.v8.3 antigen. For each reaction, 200 μg of polyclonal Fab (at 1mg/mL) was incubated with 15 μg of the antigen (at 1.4mg/mL) for 16-20 hrs (RT). Subsequently, all samples were subjected to SEC (Superose 6 Increase column, TBS as a running buffer). Fractions corresponding to the immune complexes were pooled and concentrated with Amicon ultrafiltration units (10 kDa molecular weight cutoff; MerckMillipore). Concentrated complex samples were diluted to 100 μg/mL and loaded onto the carbon-coated Cu grids (400-mesh, glow-discharged at 15 mA for 25 s prior to sample application). Filter paper was used to blot off the samples and the grids were then stained with 2 % (w/v) uranyl-formate for 60 s.

Grid imaging was performed as described above in the negative stain electron microscopy method section and using the same microscopy equipment. Appion data processing package was applied for auto-picking and particle-extraction steps^51^. 120,000 – 200,000 particles are then 2D-classified into 250 classes (50 iterations) using Relion 3.0^54^. Particles from classes with complex-like features (~60-90% of the dataset after extraction) were selected for 3D classification in Relion 3.0. Low-resolution map of non-liganded HIV Env ectodomain was used as an initial model for the 3D classification and refinement steps. For the initial 3D classification, 40 classes were used. Particles from similar looking classes were then pooled together and subjected to one or more additional rounds of 3D classification. 3D classes with unique structural features (in terms of bound polyclonal Fab) were subjected to 3D auto-refinement in Relion 3.0. 3D-refined maps were loaded into UCSF Chimera 1.13 for visualization, segmentation and figure preparation^55^. 3D refinement was also performed on the particles following the initial 2D classification step and these refined models were submitted to EMDB (1 for each dataset; see Data availability).

### Serum antibody ELISA

A 6.5 nM concentration of His-tagged 16055 SOSIP.v8.3 trimer in TBS was added to a 96-well Ni-NTA plate (Qiagen) for 2 h at RT. The plates were blocked for 30 min with TBS/2% skimmed milk. Three-fold serial dilutions of rabbit sera, starting from a 1:100 dilution (week 0,4,6,12,20) or 1:1000 dilution (week 22), were added in TBS/2% skimmed milk/20% sheep serum for a 2h incubation at RT. A 1:3000 dilution of HRP-labeled goat anti-rabbit IgG (Jackson Immunoresearch) in TBS/2% skimmed milk was added for 1 h at RT. Between each step, the plates were washed three times with TBS. Finally, after washing the plates five times with TBS/0.05% Tween-20, developing solution (1% 3,3’,5,5’-tetranethylbenzidine (Sigma-Aldrich), 0.01% H2O2, 100 mM sodium acetate and 100 mM citric acid) was added. Development of the colorimetric endpoint proceeded for 1 min before termination by adding 0.8 M H2SO4. The same procedure was used to measure binding antibody titers against the unmodified I53-50NP, except that the I53-50NP coating concentration was 3 nM and the serum starting dilution was 1:500. Binding titers (ED50-values) were determined as the dilution of serum that gave 50% of the maximal response from a sigmoidal curve.

### Competition ELISA

Half-well 96-well plates were coated overnight at RT with Galanthus nivalis lectin (Vector laboratories) at 20 μg/mL in 0.1M NaHCO3 pH 8.6. The next day plates were washed and Casein blocking buffer (ThermoScientific) was added. After 30 min a 2 μg/mL concentration of untagged 16055 SOSIP.v8.3 in Casein was added for 2 h at RT. Plates were then washed and a 1:100 dilution of rabbit sera in TBS/2% skimmed milk/20% sheep serum was added for 30 min. Next, mAbs were added for 1.5 h at RT at a final concentration that gave 80% of maximal binding signal as determined by an ELISA in the absence of serum (D11A.F9: 0.1 μg/mL; D11A.F2: 0.1 μg/mL; RM19R: 2 μg/mL, in TBS/2% skimmed milk/20% sheep serum). Plates were then washed three times and a 1:3000 dilution of HRP-labeled donkey anti-human IgG (Jackson Immunoresearch) in Casein was added for 1 h at RT. Washing and colorimetric detection then proceeded as explained for Serum antibody ELISA above. Residual binding was determined as described previously^27^.

### Neutralization assay

Neutralization assays were performed as described recently^49^. Briefly, a 1:20 dilution of heat-inactivated serum was serially diluted in 3-fold steps and mixed with Env-pseudotyped virus. After a 1 hour incubation at RT the mix was added to TZM-bl cells which were seeded to a density of 17.000 cells/well a day prior. After 72 hrs the cell medium was removed, the cells were lysed and luciferase was measured using the Bright Glo Luciferase kit (Promega). Luciferase activity was measured using a Glomax plate reader. Neutralization titers (ID50-values) were determined as the serum dilution at which infectivity was inhibited by 50%.

### Statistical analysis

Comparisons between two rabbit groups were made using a two-tailed Mann-Whitney *U* test. Correlations were analyzed by calculating Spearman’s rank correlation coefficients. Graphpad Prism 7.0 was used for statistical analyses and to determine ED50 and ID50 values for serum antibody ELISAs and neutralization assays, respectively.

## Data availability

The data supporting the findings of the study are available from the corresponding authors upon reasonable request. 3D refined models of 16066 SOSIP complexed with polyclonal Fabs were submitted to EMDB. The list of EMDB IDs: 22714 (16055 SOSIP + r2463 Poly Fab); 22715 (16055 SOSIP + r2464 Poly Fab); 22716 (16055 SOSIP + r2465 Poly Fab); 22717 (16055 SOSIP + r2466 Poly Fab); 22718 (16055 SOSIP + r2467 Poly Fab); 22719 (16055 SOSIP + r2468 Poly Fab); 22720 (16055 SOSIP + r2469 Poly Fab), 22721 (16055 SOSIP + r2470 Poly Fab); 22722 (16055 SOSIP + r2471 Poly Fab); 22723 (16055 SOSIP + r2472 Poly Fab).

## Acknowledgements

We thank Michel Nussenzweig, James Robinson, Dennis Burton, Peter Kwong, Mark Connors, John Mascola, and William Olson for donating antibodies and reagents directly or through the NIH AIDS Research and Reference Reagent Program. The following reagent was obtained through the NIH AIDS Reagent Program, Division of AIDS, NIAID, NIH: Ramos B cells from Drs. Li Wu and Vineet N. KewalRaman. We thank Andrew McGuire for kindly sharing the pRRL.EuB29 lentiviral vector that was used to transduce Ramos B cells. This work was supported by the U.S. National Institutes of Health Grant P01 AI110657 (to J.P.M., A.B.W., P.J.K., and R.W.S.) and NIAID Contract #HHSN27201100016C (to D.C.M.); by the Bill and Melinda Gates Foundation through the Collaboration for AIDS Vaccine Discovery (CAVD), grants OPP1111923 and OPP1132237 (to D.B., N.K., J.P.M., and R.W.S.), INV-008352/OPP1153692 (to M.C.), and OPP1115782 (A.B.W.); by the National Institute for Allergy and Infectious Diseases through the Scripps Consortium for HIV Vaccine Development (CHAVD) grant AI144462 (to M.C.); by the European Union’s Horizon 2020 research and innovation program under grant agreement No. 681137 (R.W.S. and M. Cr.); and by the Fondation Dormeur, Vaduz (to R.W.S.). R.W.S. is a recipient of a Vici grant from the Netherlands Organization for Scientific Research (NWO). M.J.G. is a recipient of an AMC Fellowship and a Mathilde Krim Fellowship from the American Foundation for AIDS Research (amfAR) (109514-61-RKVA). The electron microscopy data were collected at Electron Microscopy Facility of the Scripps Research Institute.

## Competing interests

N.P.K. is a co-founder, shareholder, and chair of the scientific advisory board of Icosavax, Inc. All other authors declare no competing interests.

## Author contributions

P.J.M.B., A.A, M.G., J.D.A., J.P.M., P.J.K., N.P.K., A.B.W., and R.W.S. conceived and designed experiments. P.J.M.B., A.A, M.G., J.D.A., T.P.L.B., A.Y., J.A.B., G.O., J.L.T., and R.R., performed experiments. P.J.M.B., A.A, M.G., J.D.A., G.O., C.L., D.M., J.P.M., P.J.K., M.C., N.P.K., A.B.W., and R.W.S. analyzed and interpreted data. P.J.M.B., A.A., M.G., J.D.A., J.P.M. and R.W.S. wrote the manuscript with input from all authors.

