## Supplementary Information for "Immunofocusing and enhancing autologous Tier-2 HIV-1 neutralization by displaying Env trimers on two-component protein nanoparticles"

| | Mutation | SOSIP nomenclature | | | | | Reference |
|---|---|---|---|---|---|---|---|
| | | v4 | v5 | v8.1 | v8.2 | v8.3 | |
| | 501C-605C | | | | | | Sanders et al., Plos Pathogens 2013 |
| | 559P | | | | | | |
| | R6 | | | | | | |
| | .664 | | | | | | |
| | 448N | | | | | | |
| | 64K/66R[a] | | | | | | de Taeye et al., Cell 2015; Dey et al., Virology 2008 |
| | 315Q | | | | | | |
| | 316W | | | | | | |
| | 535M[b] | | | | | | |
| | 543N/543Q[c] | | | | | | |
| | 73C-561C[d] | | | | | | Torrents de la Pena et al., Cell Reports 2017 |
| TD8 | 47D | | | | | | Guenaga et al., Journal of Virology 2016 |
| | 49E | | | | | | |
| | 65K | | | | | | |
| | 106T | | | | | | |
| | 165L | | | | | | |
| | 429R | | | | | | |
| | 432Q | | | | | | |
| | 500R | | | | | | |
| MD39 | 106E | | | | | | Steichen et al., Immunity 2017 |
| | 271I | | | | | | |
| | 288L | | | | | | |
| | 304V | | | | | | |
| | 319Y | | | | | | |
| | 363Q[e] | | | | | | |
| | 519S | | | | | | |
| | 561P[f] | | | | | | |
| | 568D | | | | | | |
| | 570H | | | | | | |
| | 585H | | | | | | |
| | 569G | | | | | | Guenaga et al., Immunity 2017 |

**Supplementary Table 1.** Overview of mutations that make up SOSIP v4-v8.3.

[a] The variant 64K (v4.1) is used for BG505 and B41; while the variant 66R (v4.2) is used for the rest of strains.

[b] This mutation was not introduced in 16055 SOSIP.

[c] de Taeye et al. identified two variants (543N and 543Q) with positive effects. 16055 naturally contains a Q at position 543.

[d] 72C-564C (v5.1) is an alternative disulfide bond that works as well as 73C-561C (v5.2), but we generally work with the latter.

[e] Is left out in several genotypes (BG505) as it introduces a glycan hole. This is not the case for 16055.

[f] Mutation 561P (Steichen et al, Immunity 2017) is not included, as we routinely introduce a Cys in this position for the v5 disulfide bond.

### 16055 SOSIP

| | 88 | 160 | 187 | 197 | 262 | 276 | 289 | 301 | 360 | 386 | 409 | 442 | 448 | 463 | 611 | 616 | 625 | 637 |
|---|---|---|---|---|---|---|---|---|---|---|---|---|---|---|---|---|---|---|
| High Mannose | 41 | 65 | 56 | 34 | 100 | 90 | 100 | 62 | 100 | 100 | 49 | 93 | 100 | 43 | 0 | 1 | 35 | 84 |
| M9 | 0 | 1 | 0 | 0 | 83 | 0 | 68 | 0 | 68 | 78 | 0 | 17 | 5 | 0 | 0 | 0 | 0 | 0 |
| M8 | 0 | 11 | 0 | 13 | 14 | 7 | 28 | 0 | 20 | 20 | 0 | 67 | 67 | 0 | 0 | 0 | 0 | 0 |
| M7 | 0 | 3 | 0 | 13 | 1 | 1 | 4 | 0 | 10 | 2 | 0 | 1 | 14 | 0 | 0 | 1 | 5 | 17 |
| M6 | 8 | 19 | 0 | 4 | 1 | 43 | 0 | 0 | 1 | 0 | 0 | 4 | 8 | 0 | 0 | 0 | 0 | 24 |
| M5 | 27 | 28 | 54 | 0 | 0 | 33 | 0 | 62 | 0 | 0 | 49 | 3 | 5 | 39 | 0 | 0 | 29 | 32 |
| M4 | 0 | 1 | 0 | 0 | 0 | 1 | 0 | 0 | 1 | 0 | 0 | 0 | 1 | 0 | 0 | 0 | 0 | 3 |
| M3 | 0 | 0 | 0 | 0 | 0 | 0 | 0 | 0 | 0 | 0 | 0 | 0 | 0 | 0 | 0 | 0 | 0 | 0 |
| Hybrid | 5 | 1 | 0 | 2 | 0 | 2 | 0 | 0 | 0 | 0 | 0 | 0 | 0 | 2 | 0 | 0 | 0 | 7 |
| FHybrid | 0 | 0 | 2 | 1 | 0 | 0 | 0 | 0 | 0 | 0 | 0 | 0 | 0 | 2 | 0 | 0 | 0 | 0 |
| HexNAC | 1 | 0 | 0 | 1 | 0 | 1 | 0 | 0 | 0 | 0 | 0 | 0 | 0 | 1 | 0 | 0 | 0 | 1 |
| A1 | 10 | 1 | 2 | 0 | 0 | 2 | 0 | 0 | 0 | 0 | 0 | 0 | 0 | 0 | | | 4 | 6 |
| FA1 | 0 | 0 | 2 | 7 | 0 | 0 | 0 | 35 | 0 | 0 | 39 | 0 | 0 | 10 | | | 3 | 3 |
| A2/A1B | 1 | 0 | 1 | 0 | 0 | 1 | 0 | 0 | 0 | 0 | 0 | 0 | 0 | 0 | | | 7 | 0 |
| FA2/FA1B | 17 | 1 | 36 | 3 | 0 | 0 | 0 | 0 | 0 | 0 | 13 | 0 | 0 | 41 | 0 | 0 | 36 | 2 |
| A3/A2B | 4 | 0 | 0 | 0 | 0 | 0 | 0 | 0 | 0 | 0 | 0 | 0 | 0 | 0 | | | 0 | 0 |
| FA3/FA2B | 19 | 0 | 3 | 0 | 0 | 0 | 0 | 0 | 0 | 0 | 0 | 0 | 0 | 4 | | | 11 | 2 |
| A4/A3B | 0 | 0 | 0 | 0 | 0 | 0 | 0 | 0 | 0 | 0 | 0 | 0 | 0 | 0 | | | 0 | 0 |
| FA4/FA3B | 0 | 0 | 0 | 0 | 0 | 0 | 0 | 0 | 0 | 0 | 0 | 0 | 0 | 1 | | | 0 | 0 |
| Unoccupied | 6 | 34 | 1 | 56 | 0 | 7 | 0 | 3 | 0 | 0 | 0 | 7 | 0 | 0 | 0 | 60 | 3 | 2 |

| | 88 | 136 | 139 | 145 | 156 | 160 | 187 | 197 | 230 | 234 | 241 | 262 | 276 | 289 | 301 | 360 | 386 | 392 | 399 | 404 | 409 | 442 | 448 | 463 | 611 | 616 | 625 | 637 |
|---|---|---|---|---|---|---|---|---|---|---|---|---|---|---|---|---|---|---|---|---|---|---|---|---|---|---|---|---|
| High Mannose | 42 | 69 | 69 | 0 | 63 | 65 | 56 | 34 | 35 | 100 | 94 | 100 | 90 | 100 | 62 | 100 | 100 | | | 0 | 49 | 93 | 100 | 43 | 8 | 1 | 35 | 84 |
| Complex | 52 | 18 | 31 | 100 | 1 | 2 | 43 | 11 | 0 | 0 | 0 | 0 | 3 | 0 | 35 | 0 | 0 | | | 100 | 51 | 0 | 0 | 57 | 63 | 39 | 62 | 14 |
| Unoccupied | 6 | 14 | 0 | 0 | 36 | 34 | 1 | 56 | 65 | 0 | 6 | 0 | 7 | 0 | 3 | 0 | 0 | | | 0 | 0 | 7 | 0 | 0 | 29 | 60 | 3 | 2 |

### 16055 SOSIP-I53-50A

| | 88 | 145 | 156 | 160 | 187 | 197 | 234 | 241 | 262 | 276 | 289 | 301 | 360 | 386 | 409 | 442 | 448 | 463 | 625 | 637 |
|---|---|---|---|---|---|---|---|---|---|---|---|---|---|---|---|---|---|---|---|---|
| High Mannose | 69 | 58 | 44 | 84 | 56 | 66 | 100 | 0 | 100 | 97 | 100 | 82 | 99 | 100 | 42 | 97 | 98 | 16 | 86 | 93 |
| M9 | 0 | 0 | 23 | 0 | 0 | 0 | 76 | 100 | 89 | 1 | 18 | 7 | 74 | 80 | 0 | 73 | 0 | 0 | 0 | 0 |
| M8 | 0 | 0 | 17 | 0 | 0 | 25 | 24 | 0 | 11 | 13 | 58 | 0 | 20 | 18 | 0 | 4 | 87 | 0 | 0 | 17 |
| M7 | 0 | 4 | 0 | 12 | 0 | 12 | 0 | 0 | 0 | 11 | 25 | 0 | 1 | 2 | 7 | 0 | 0 | 0 | 1 | 40 |
| M6 | 30 | 0 | 1 | 29 | 13 | 0 | 0 | 0 | 1 | 43 | 0 | 6 | 0 | 1 | 2 | 7 | 7 | 9 | 0 | 5 |
| M5 | 33 | 46 | 3 | 39 | 41 | 17 | 0 | 0 | 0 | 26 | 0 | 64 | 1 | 0 | 33 | 11 | 2 | 1 | 84 | 28 |
| M4 | 1 | 0 | 0 | 0 | 1 | 0 | 0 | 0 | 0 | 1 | 0 | 0 | 0 | 0 | 0 | 0 | 1 | 0 | 0 | 1 |
| M3 | 0 | 0 | 0 | 0 | 0 | 0 | 0 | 0 | 0 | 0 | 0 | 0 | 0 | 0 | 0 | 0 | 0 | 0 | 0 | 0 |
| Hybrid | 3 | 0 | 0 | 2 | 0 | 6 | 0 | 0 | 0 | 0 | 0 | 0 | 1 | 0 | 0 | 0 | 0 | 0 | 1 | 0 |
| FHybrid | 0 | 0 | 0 | 0 | 0 | 0 | 0 | 0 | 0 | 0 | 0 | 0 | 0 | 0 | 0 | 0 | 0 | 1 | 0 | 0 |
| HexNAc | 2 | 9 | 0 | 2 | 1 | 6 | 0 | 0 | 0 | 1 | 0 | 5 | 1 | 0 | 0 | 2 | 0 | 5 | 0 | 3 |
| A1 | 8 | 0 | 0 | 1 | 3 | 0 | 0 | 0 | 0 | 2 | 0 | 0 | 0 | 0 | 0 | 0 | 2 | 4 | 0 | 0 |
| FA1 | 0 | 5 | 0 | 0 | 0 | 0 | 0 | 0 | 0 | 0 | 0 | 0 | 0 | 0 | 0 | 1 | 0 | 40 | 0 | 2 |
| A2/A1B | 12 | 0 | 0 | 0 | 0 | 1 | 0 | 0 | 0 | 0 | 0 | 0 | 0 | 0 | 0 | 0 | 0 | 5 | 7 | 0 |
| FA2/FA1B | 0 | 4 | 0 | 0 | 40 | 13 | 0 | 0 | 0 | 0 | 0 | 0 | 0 | 0 | 0 | 0 | 0 | 24 | 1 | 4 |
| A3/A2B | 3 | 0 | 0 | 0 | 0 | 0 | 0 | 0 | 0 | 0 | 0 | 0 | 0 | 0 | 0 | 0 | 0 | 0 | 1 | 0 |
| FA3/FA2B | 7 | 0 | 0 | 0 | 1 | 3 | 0 | 0 | 0 | 0 | 0 | 0 | 0 | 0 | 20 | 0 | 0 | 8 | 4 | 1 |
| A4/A3B | 0 | 0 | 0 | 0 | 0 | 0 | 0 | 0 | 0 | 0 | 0 | 0 | 0 | 0 | 0 | 0 | 0 | 0 | 0 | 0 |
| FA4/FA3B | 0 | 0 | 0 | 0 | 0 | 0 | 0 | 0 | 0 | 0 | 0 | 0 | 0 | 0 | 0 | 0 | 0 | 4 | 0 | 0 |
| Unoccupied | 0 | 32 | 56 | 15 | 0 | 17 | 0 | 0 | 0 | 1 | 0 | 18 | 1 | 0 | 37 | 2 | 0 | 0 | 0 | 0 |

| | 88 | 136 | 139 | 145 | 156 | 160 | 187 | 197 | 230 | 234 | 241 | 262 | 276 | 289 | 301 | 360 | 386 | 392 | 399 | 404 | 409 | 442 | 448 | 463 | 611 | 616 | 625 | 637 |
|---|---|---|---|---|---|---|---|---|---|---|---|---|---|---|---|---|---|---|---|---|---|---|---|---|---|---|---|---|
| High Mannose | 69 | 97 | 93 | 58 | 44 | 84 | 56 | 66 | n.d. | 100 | 100 | 100 | 97 | 100 | 82 | 99 | 100 | n.d. | n.d. | 0 | 42 | 97 | 98 | 16 | 41 | 14 | 86 | 93 |
| Complex | 30 | 0 | 0 | 10 | 0 | 1 | 44 | 17 | n.d. | 0 | 0 | 0 | 3 | 0 | 0 | 0 | 0 | n.d. | n.d. | 100 | 20 | 2 | 2 | 84 | 42 | 44 | 13 | 7 |
| Unoccupied | 0 | 3 | 7 | 32 | 56 | 15 | 0 | 17 | n.d. | 0 | 0 | 0 | 1 | 0 | 18 | 1 | 0 | n.d. | n.d. | 0 | 37 | 2 | 0 | 0 | 16 | 42 | 0 | 0 |

**Supplementary Table 2.** Glycan composition of 16055 SOSIP and 16055 SOSIP-I53-50A. Glycoforms are shown for each PNGS of 16055 SOSIP (top) and 16055 SOSIP-I53-50A (bottom), with each number representing the percentage of the specific glycoform as stated in the left column. Oligomannos/hybrid-type glycans are shaded in green and complex glycans are shaded in pink. The absence of a glycan is shaded in grey. Glycoforms are categorized according to the number of mannose residues (M3-M9), hybrid glycans by the precense/absence of fucose (FHybrid and Hybrid), and complex glycans by the presence/absence of fucose and the number of antenna. The upper table for each construct contains a breakdown of compositions obtained using intact glycopeptides. The percentages in lower table are obtained by using glycosidase-treated peptides (see Methods for details). Sites that appear in the lower table but not the upper are those that could not be obtained using intact glycopeptides. n.d. = not determined.

#### Week 6

| Immunogen | Rabbit ID | Control | | Autologous | |
|---|---|---|---|---|---|
| | Virus | MLV | | 16055 | |
| | Tier | - | | 2 | |
| | Clade | - | | C | |
| | Lab | AMC | DUMC | AMC | DUMC |
| 16055 SOSIP | 2463 | 27 | n.d. | 20 | n.d. |
| | 2464 | 21 | n.d. | <20 | n.d. |
| | 2465 | 54 | n.d. | 92 | n.d. |
| | 2466 | <20 | n.d. | <20 | n.d. |
| | 2467 | <20 | n.d. | <20 | n.d. |
| 16055 SOSIP-I53-50NP | 2468 | <20 | n.d. | 87 | n.d. |
| | 2469 | 22 | n.d. | 55 | n.d. |
| | 2470 | <20 | n.d. | <20 | n.d. |
| | 2471 | 20 | n.d. | 92 | n.d. |
| | 2472 | 22 | n.d. | 26 | n.d. |

#### Week 22

| Immunogen | Rabbit ID | Control | | Autologous | | Heterologous | |
|---|---|---|---|---|---|---|---|
| | Virus | MLV | | 16055 | | SF162 | MW965 |
| | Tier | - | | 2 | | 1A | 1A |
| | Clade | - | | C | | B | C |
| | Lab | AMC | DUMC | AMC | DUMC | AMC | DUMC |
| 16055 SOSIP | 2463 | <20 | <20 | 20 | 20 | 123 | 1672 |
| | 2464 | <20 | <20 | 2814 | 1793 | <20 | 837 |
| | 2465 | <20 | <20 | 376 | 158 | <20 | 348 |
| | 2466 | <20 | <20 | 60 | 25 | <20 | 108 |
| | 2467 | <20 | <20 | 474 | 186 | <20 | 212 |
| 16055 SOSIP-I53-50NP | 2468 | <20 | <20 | 18660 | 4870 | <20 | 479 |
| | 2469 | <20 | <20 | 13348 | 4333 | <20 | 109 |
| | 2470 | <20 | <20 | 8363 | 2923 | <20 | 3585 |
| | 2471 | <20 | <20 | 67677 | 26174 | <20 | 2000 |
| | 2472 | <20 | <20 | 4200 | 1550 | <20 | 563 |

**Supplementary Table 3.** Midpoint neutralization titers at week 6 and 22 from rabbits that received 16055 SOSIP or 16055 SOSIP-I53-50NP tested against a panel of Env-pseudoviruses. TZM-bl neutralization assays were performed either at the AMC and/or DUMC as indicated above each column. $ID_{50}$ values, i.e. the serum dilution at which infectivity was inhibited by 50%, are shown and color coded: white = no neutralization, $ID_{50}$ < 20; grey = very weak neutralization, 20 > $ID_{50}$ >40; yellow = weak neutralization, 40 > $ID_{50}$ > 100; orange = moderate neutralization, 100 > $ID_{50}$ > 1000; red = strong neutralization, 1000 > $ID_{50}$ > 10000; purple = very strong neutralization, $ID_{50}$ > 10000. MLV = murine leukemia virus (negative control). n.d. = not determined.

#### Week 22

| Immunogen | Rabbit ID | Heterologous | | | | | | | | |
|---|---|---|---|---|---|---|---|---|---|---|
| | Virus | 25710-2.43 | TRO.11 | BJOX002000.03.2 | Ce1176_A3 | X1632_S2_B10 | 246-F3_C10_2 | CH119.10 | Ce703010217_B6 | CNE55 |
| | Tier | 2 | 2 | 2 | 2 | 2 | 2 | 2 | 2 | 2 |
| | Clade | | | | | | | | | |
| | Lab | DUMC | DUMC | DUMC | DUMC | DUMC | DUMC | DUMC | DUMC | DUMC |
| 16055 SOSIP | 2463 | <20 | <20 | <20 | <20 | <20 | <20 | <20 | <20 | <20 |
| | 2464 | <20 | <20 | <20 | <20 | <20 | <20 | 25 | <20 | <20 |
| | 2465 | <20 | <20 | <20 | 51 | <20 | 28 | 44 | <20 | 21 |
| | 2466 | <20 | <20 | <20 | <20 | <20 | <20 | 21 | <20 | <20 |
| | 2467 | <20 | <20 | <20 | <20 | <20 | <20 | <20 | <20 | <20 |
| 16055 SOSIP-I53-50NP | 2468 | <20 | <20 | <20 | 28 | <20 | <20 | 34 | <20 | <20 |
| | 2469 | <20 | <20 | <20 | <20 | <20 | <20 | <20 | <20 | <20 |
| | 2470 | <20 | <20 | <20 | <20 | <20 | <20 | <20 | <20 | <20 |
| | 2471 | <20 | <20 | <20 | <20 | <20 | <20 | <20 | <20 | <20 |
| | 2472 | <20 | <20 | <20 | 40 | <20 | <20 | 24 | <20 | <20 |

**Supplementary Table 4.** Midpoint neutralization titers at week 22 from rabbits that received 16055 SOSIP or 16055 SOSIP-I53-50NP tested against a panel of heterologous Env-pseudoviruses. TZM-bl neutralization assays were performed at DUMC as indicated above each column. $ID_{50}$ values, i.e. the serum dilution at which infectivity was inhibited by 50%, are shown and color coded as shown in the legend for Supplementary Table 3.

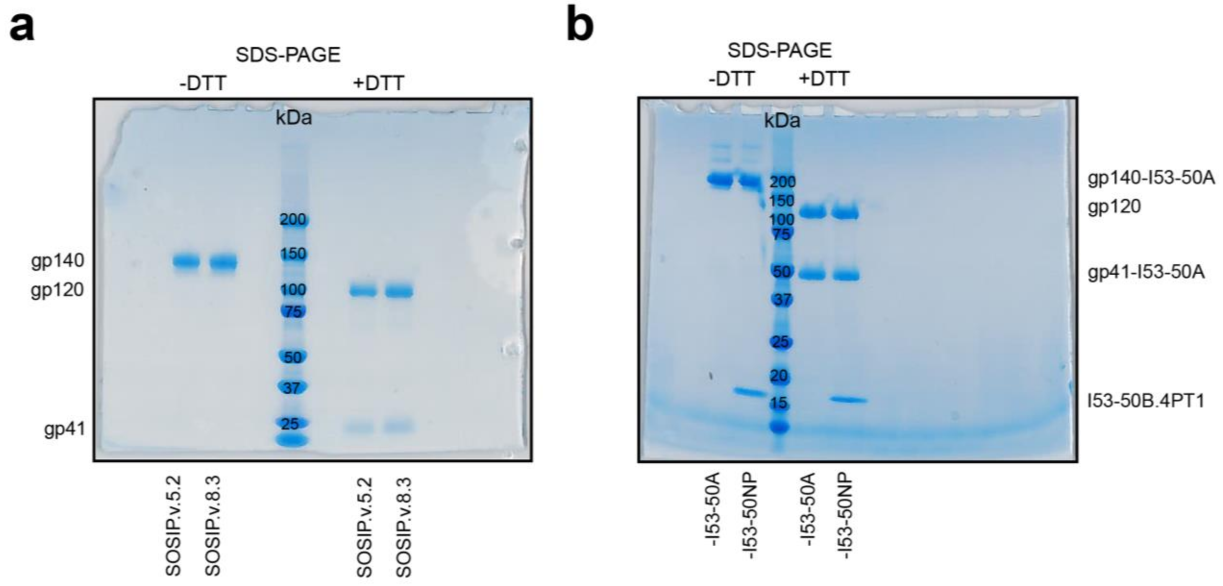


**Supplementary Fig. 1.** Biophysical characterization of 16055 SOSIPs, 16055 SOSIP-I53-50A, and 16055 SOSIP-I53-50NP **a** Non-reducing (-DTT) and reducing (+DTT) SDS-PAGE of 16055 SOSIP.v5.2 and 16055 SOSIP.v8.3. **b** As panel-a but for 16055 SOSIP-I53-50A and 16055 SOSIP-I53-50NP.

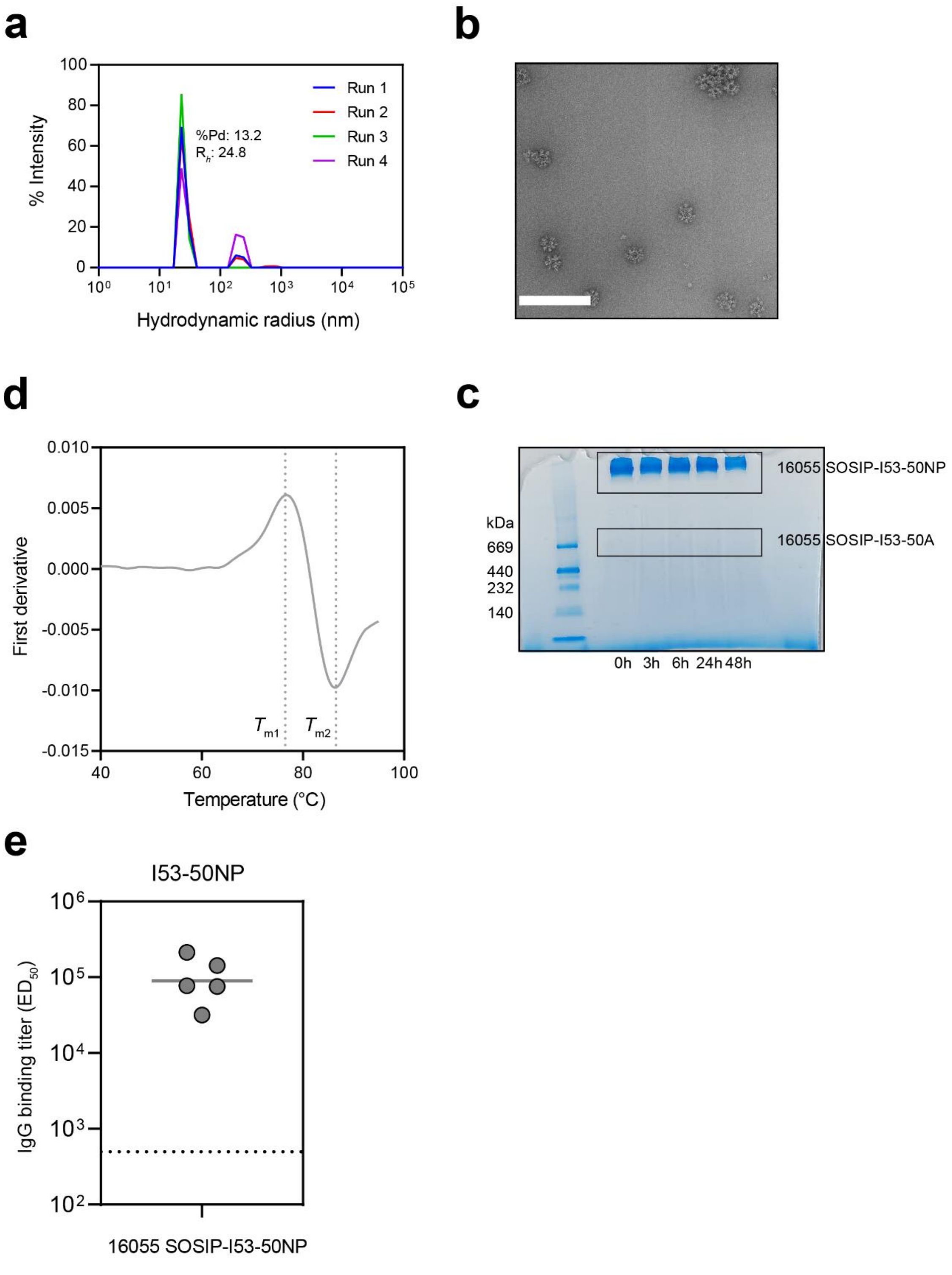


**Supplementary Fig. 2.** Biophysical characterization of 16055 SOSIP-I53-50NPs and the immunogenicity of the I53-50NP core. **a** Dynamic light scattering data of 16055 SOSIP-I53-50NPs. Average polydispersity (%Pd) and hydrodynamic radius ($R_h$) are shown for the non-aggregated 16055 SOSIP-I53-50NP population. A %Pd < 15 is considered a monodisperse population. An overlay of four individual runs is shown. **b** Raw nsEM micrograph of 16055 SOSIP-

I53-50NPs showing the small percentage of aggregates. **c** Thermostability of 16055 SOSIP-I53-50NP as determined by NanoDSF. The dotted lines indicate the melting temperatures of the 16055 SOSIP ($T_{m1}$) and I53-50NP core ($T_{m2}$). Representative melting curve of three technical replicates is shown. **d** Native-PAGE of 16055 SOSIP-I53-50NP incubated at 37°C in PBS for 0, 3, 6, 24 and 48 h. The upper box indicates the band for 16055 SOSIP-I53-50NPs whereas the lower indicates where a 16055 SOSIP-I53-50A band would have been visible upon nanoparticle disassembly. **e** Midpoint binding titers against I53-50NPs from week 22 rabbits sera of 16055 SOSIP-I35-50NP recipients. Horizontal bar indicates the geometric mean. The dotted line indicates the lowest dilution.

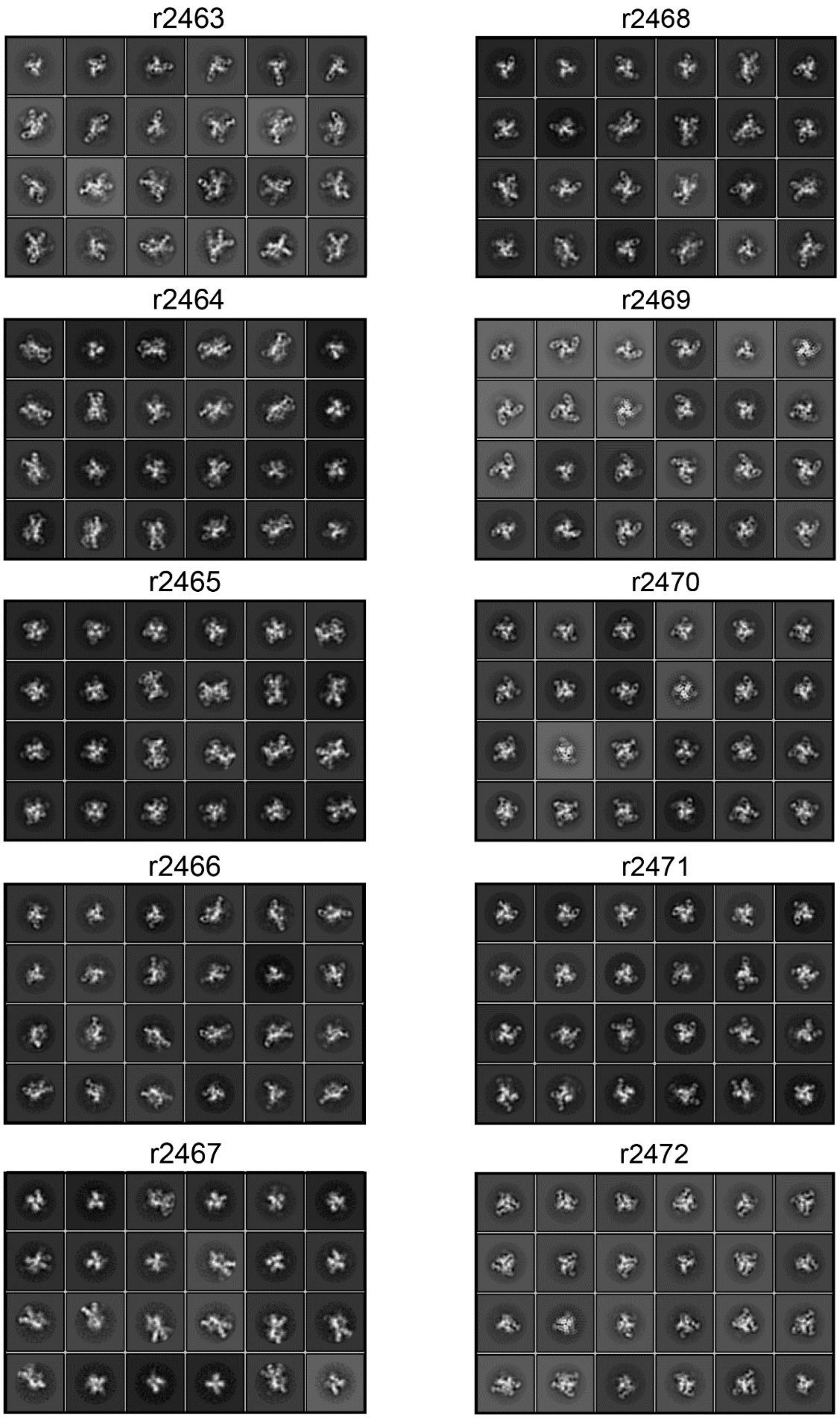


**Supplementary Fig.3.** Representative 2D class averages of 16055 SOSIP trimers complexed with polyclonal Fabs from rabbit sera with IDs indicated above.

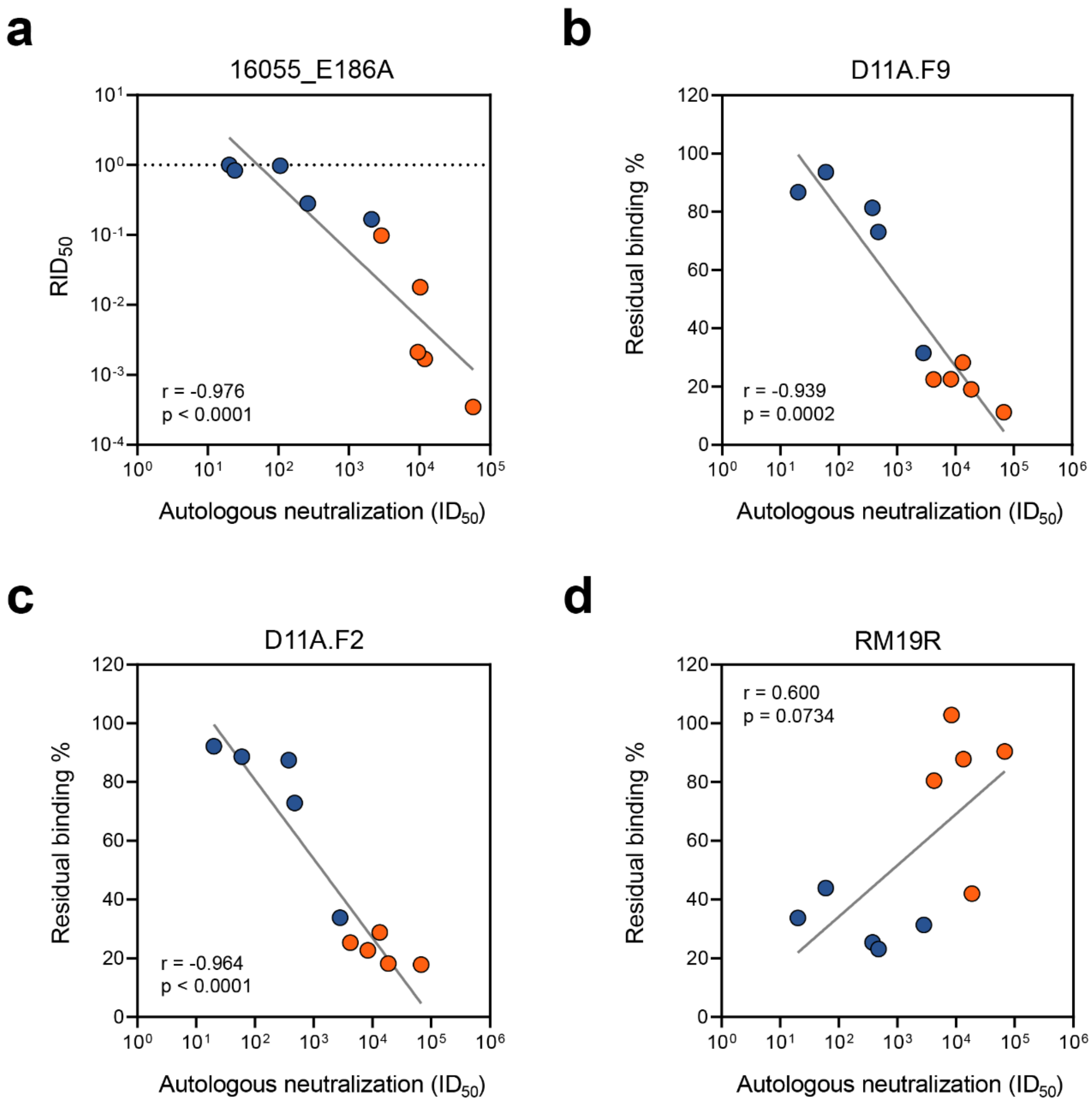


**Supplementary Fig. 4.** Epitope mapping of NAb responses in rabbits immunized with 16055 SOSIP or 16055 SOSIP-I53-50NPs. **a** Correlation plot of RID50 versus 16055 midpoint neutralization titers. **b** Correlation plot of D11A.F9 competition versus 16055 midpoint neutralization titers. **c** The same as b but then for D11A.F2 **d** The same as b but then for RM19R. **a-d** The *r* and *p* value is shown for non-parametirc Spearman rank correlation. Blue and orange symbols represent individual rabbits immunized with 16055 SOSIP or 16055 SOSIP-I53-50NPs, respectively.
